# Activation of the imprinted Prader-Willi Syndrome locus by CRISPR-based epigenome editing

**DOI:** 10.1101/2024.03.03.583177

**Authors:** Dahlia Rohm, Joshua B. Black, Sean R. McCutcheon, Alejandro Barrera, Daniel J. Morone, Xander Nuttle, Celine E. de Esch, Derek J.C. Tai, Michael E. Talkowski, Nahid Iglesias, Charles A. Gersbach

## Abstract

Epigenome editing with DNA-targeting technologies such as CRISPR-dCas9 can be used to dissect gene regulatory mechanisms and potentially treat associated disorders. For example, Prader-Willi Syndrome (PWS) is caused by loss of paternally expressed imprinted genes on chromosome 15q11.2-q13.3, although the maternal allele is intact but epigenetically silenced. Using CRISPR repression and activation screens in human induced pluripotent stem cells (iPSCs), we identified genomic elements that control expression of the PWS gene *SNRPN* from the paternal and maternal chromosomes. We showed that either targeted transcriptional activation or DNA demethylation can activate the silenced maternal *SNRPN* and downstream PWS transcripts. However, these two approaches function at unique regions, preferentially activating different transcript variants and involving distinct epigenetic reprogramming mechanisms. Remarkably, transient expression of the targeted demethylase leads to stable, long-term maternal *SNRPN* expression in PWS iPSCs. This work uncovers targeted epigenetic manipulations to reprogram a disease-associated imprinted locus and suggests possible therapeutic interventions.

## Introduction

Complex mechanisms of epigenetic regulation, such as imprinting and X-inactivation, are critical in mammalian development (Jaenisch and Bird, 2003, Ferguson-Smith and Surani, 2001, Ferguson-Smith, 2011), and dysregulation of these processes is the basis for several human disorders (Eggermann et al., 2015). Although the identity and dynamic control of the epigenetic marks involved in these processes have been investigated extensively (Patrat et al., 2020, Barlow and Bartolomei, 2014), the mechanisms of heritable and allele-specific changes in gene expression at specific loci are still relatively poorly understood. Altering specific epigenetic modifications at imprinted loci could provide insight into the contributions of these marks to the stability of gene expression patterns. The advent of epigenome editing with DNA-targeting technologies such as CRISPR-Cas9 has revolutionized our ability to manipulate gene expression states in living cells and organisms. In particular, nuclease-deactivated Cas9 (dCas9) fused to transcriptional regulators or epigenome modifiers can recruit transcription factors to promoters or enhancers, directly alter histone marks or DNA methylation, or sterically block transcription or transcription factor binding (Hilton et al., 2015, Liu et al., 2016, Shariati et al., 2019, Cano-Rodriguez et al., 2016).

dCas9-based epigenome editing can deposit chromatin modifications in a highly specific manner (Nakamura et al., 2021, Thakore et al., 2016), enabling precise evaluation of the link between chromatin modification and gene expression (Cano-Rodriguez et al., 2016, O’Geen et al., 2019, O’Geen et al., 2017, Nuñez et al., 2021). These technologies are also used for unbiased identification of distal gene regulatory elements through pooled gRNA screens (Fulco et al., 2016, Klann et al., 2017, Klann et al., 2018, Montalbano et al., 2017, Simeonov et al., 2017), defining key regulatory elements that are otherwise challenging to predict (Canver et al., 2015, Diao et al., 2016, Korkmaz et al., 2016, Sanjana et al., 2016). In addition to mapping the regulatory landscape in the human genome, these studies serve to identify potential drug targets for next-generation epigenetic therapies. We sought to apply these tools to identify regulatory elements of the imprinted 15q11-13 locus that is implicated in PWS.

PWS is a neuroendocrine and neurobehavioral disorder caused by the absence of expression of the paternal copy of imprinted genes in the chromosomal region 15q11-13 (Buiting, 2010). It is characterized clinically by hyperphagia, early-onset obesity and intellectual disability (Cassidy et al., 2012). While the exact genetic basis that causes PWS remains unclear, patient mutation profiles have implicated the small nucleolar RNA (snoRNA) gene cluster *SNORD116* downstream of the *SNURF-SNRPN* open reading frame within 15q11-13 as a likely primary contributor to the disease etiology (Li et al., 2016, Sutcliffe et al., 1994). Because the maternal copy of chromosome 15q11-13 remains intact but epigenetically silenced, activation of the maternal allele provides a therapeutic opportunity to restore expression of PWS-associated genes.

Activation of PWS-associated genes from the maternal allele has been realized through inhibition of epigenetic modifying enzymes and transcription factors (Cruvinel et al., 2014, Kim et al., 2017, Langouet et al., 2018, Saitoh and Wada, 2000). However, small molecule-based inhibition of epigenetic modifiers carries the risk of off-target effects due to the global loss of critical gene regulation enzymes, given their widespread expression and association with numerous loci (Bernstein et al., 2010, Consortium, 2012). Recently developed epigenome editing technologies offer a more targeted approach to achieve epigenetic modification at the intended locus with minimal off-target activity (Thakore et al., 2016, Nakamura et al., 2021, Chavez et al., 2023).

Here, we applied a CRISPR/dCas9-based screening approach to identify genomic regulatory elements at the 15q11-13 locus controlling expression of the *SNURF-SNRPN* host transcript in human iPSCs. By conducting independent screens against the paternal and maternal alleles, we identified regulatory elements controlling expression of PWS-associated genes. Importantly, we activated the maternal host transcript and other PWS genes using targeted epigenetic editing with different dCas9-based effectors, employing both DNA methylation-dependent and -independent mechanisms. Our findings complement previous studies using small-molecule inhibitors of epigenetic enzymes to reactivate imprinted genes and provide a promising avenue for targeted epigenetic therapy.

## Results

### Generation of *SNRPN-2A-GFP* reporter cell lines for CRISPR screening

The chromosome 15q11-13 region contains several imprinted genes. Most protein-coding genes, such as *MAGEL2*, *NDN* and *SNURF-SNRPN* (henceforth referred to only as *SNRPN*), along with numerous noncoding RNAs (ncRNAs), including snoRNA clusters *SNORD115* and *SNORD116*, are exclusively expressed from the paternally inherited allele (Figure 1A). Common genotypes in PWS patients consist of deletions within 15q11-13, affecting multiple coding and noncoding genes, with a subset of genotypes emphasizing the significance of snoRNA clusters in PWS etiology (Bieth et al., 2015, de Smith et al., 2009, Duker et al., 2010, Sahoo et al., 2008). *SNURF-SNRPN* and downstream ncRNAs, including *SPA* RNAs and snoRNAs, are processed from the single host transcript *SNHG14* that initiates upstream of the PWS-IC (Wu et al., 2016, Chung et al., 2020). Notably, the canonical *SNRPN* promoter and exon 1 regions overlap the PWS-IC, but some transcript variants initiate upstream of the PWS-IC and include alternative noncoding exons of *SNRPN* located upstream of the PWS-IC (referred to as “upstream exons”) and expressed in neurons (Dittrich et al., 1996, Landers et al., 2004, Langouet et al., 2018). Considering the role of the PWS-IC in imprinting within 15q11-13, we selected *SNRPN* expression as a proxy for the imprinting status of the PWS locus.

**Figure 1.**
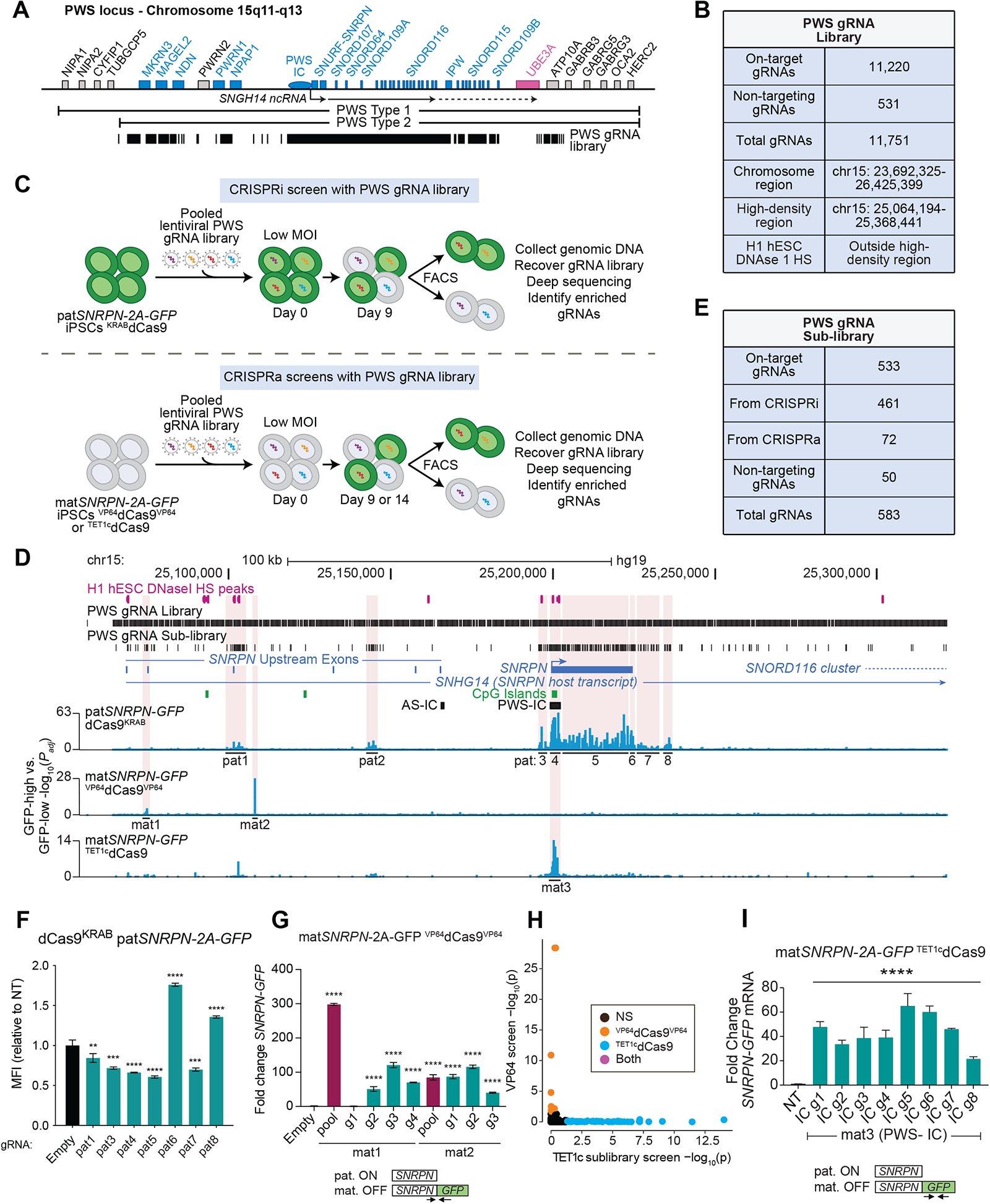
High-throughput screens reveal regulatory elements of maternal and paternal *SNRPN* alleles. (A) Schematic of the PWS locus on chr15 with common PWS deletions and the PWS gRNA library. Each thin vertical line represents a single gRNA. Genes colored blue are maternally imprinted, those that are pink are paternally imprinted, and those that are grey are not imprinted. (B) Summary of the PWS gRNA library. (C) Schematic of experimental protocol for CRISPRa/CRISPRi screens. (D) CRISPR screen results (zoomed in, see Supp. Fig. S1C) displayed as -log_10_(p_ad_j), where p_ad_j is the multiple-hypothesis-corrected p-value from DESeq2. Notable regions are highlighted in beige. (E) Summary of the PWS gRNA sub-library. (F) Flow cytometry of *SNRPN-GFP* MFI for validations of individual gRNAs of the pat*SNRPN-2A-GFP* CRISPRi dCas9^KRAB^ screen. MFI values normalized to Empty vector. One-way ANOVA followed by Dunnett’s multiple comparisons test vs. Empty. **p < 0.01, ***p < 0.001, ****p<0.0001. (G) qPCR of *SNRPN-GFP* from individual gRNA validations of each of the gRNAs in the mat1 and mat2 pools shown in Fig. 1F. (H) Plot of -log_10_(p_adj_) values of each gRNA in the ^VP64^dCas9^VP64^ full library screen vs. ^Tet1c^dCas9 sublibrary screen, plotting only the gRNAs present in both screens. (Significant p_adj_ < 0.05.) (I) qPCR of *SNRPN-GFP* for validations of individual gRNAs of the mat*SNRPN-2A-GFP* CRISPRa ^Tet1c^dCas9 screen. For qPCR in (G) and (I), fold change values are plotted mean +/-SD, but statistics were calculated on ddCt values (normalized to *GAPDH* and empty or NT vector sample); one-way ANOVA followed by Dunnett’s test vs. empty vector. ***p<0.001, ****p<0.0001. Unmarked comparisons are not significant.

Through CRISPR/Cas9-based homologous recombination, we inserted the coding sequence of superfolder GFP (sfGFP) into exon 10 of *SNRPN* in a wild-type human iPSC line (Supp. Figure S1A,B). To ensure proper functionality, localization, and stability of SNRPN, we included a P2A skipping peptide between SNRPN and sfGFP, avoiding disruption that could result from direct fusion to GFP. We chose a wild-type iPSC line with two intact copies of 15q11-13 in order to isolate heterozygous clones with *SNRPN-2A-GFP* on either the maternal or paternal allele. GFP-positive cells were expected exclusively in the paternally tagged cells if imprinting was maintained (Supp. Figure S1A). We derived clonal lines with either allele tagged, and heterozygous clones displayed the expected bimodal distribution in GFP fluorescence (Supp. Figure S1B). We selected two *SNRPN-2A-GFP* lines that independently report on *SNRPN* expression from the paternal or maternal allele. These cell lines were used in all subsequent CRISPR screens described in this study.

### Identification of allele-specific regulatory elements with CRISPRa and CRISPRi screens in *SNRPN-2A-GFP* hiPSCs

Epigenetic marks within the 15q11-13 region that are specific to the parent-of-origin are associated with allele-specific expression of PWS genes (Horsthemke and Wagstaff, 2008). Furthermore, putative cis-acting regulatory sequences have been identified within the PWS locus (Horsthemke and Wagstaff, 2008). We reasoned that CRISPR-based epigenetic editing could identify and reveal the function of regulatory regions that control the maintenance of imprinting at this locus. Therefore, we designed a gRNA library within the 15q11-13 locus to screen for regulatory elements controlling expression of paternal or maternal *SNRPN-2A-GFP* (Fig. 1A,B, Supp. Figure S1C).

Previous studies using CRISPR-based screening approaches to uncover regulatory elements often found that these elements were located in the proximity (i.e. within a megabase) of their target genes and often were annotated with canonical markers of regulatory activity, such as DNase I hypersensitivity (Klann et al., 2018, Montalbano et al., 2017). However, it is possible that elements that establish imprinting at early stages of differentiation and development no longer harbor canonical signatures of regulatory elements, and thus perturbing their function would require unbiased screening with gRNA libraries tiling the region. Consequently, our designed gRNA library consisted of a high-density region covering ∼300 kilobases (kb) centered at the PWS-IC and extending upstream to alternative *SNRPN* exons and downstream to the *SNORD116* cluster. Additional gRNAs outside of the high-density region were designed to target putative regulatory elements throughout the remaining imprinted region based on DNase I hypersensitivity signal in human embryonic stem cells (Consortium, 2012). The full gRNA library consisted of 11,751 total gRNAs, including 531 non-targeting controls (Figure 1B, and Supplementary Table 1).

We performed two independent screens in the paternal and maternal *SNRPN-2A-GFP* cell lines (Figure 1C). A CRISPRi screen with the dCas9^KRAB^ repressor in paternally tagged *SNRPN-2A-GFP* cells (*patSNRPN-2A-GFP*) was performed to identify regions that can coordinate repression of *SNRPN* transcription. dCas9^KRAB^ is widely used for targeted gene repression across various epigenetic contexts, including promoters and distal regulatory elements (Gilbert et al., 2014, Gilbert et al., 2013, Thakore et al., 2015). Similarly, we performed a CRISPRa screen in the maternally tagged *SNRPN-2A-GFP* cells (*matSNRPN-2A-GFP*) using the ^VP64^dCas9^VP64^ activator. The choice of the VP64 transactivation domain was based on its reported broad activity across diverse chromatin contexts and its ability to initiate chromatin remodeling at the target site (Gao et al., 2013, Polstein et al., 2015). The double fusion of VP64 to both termini of dCas9 (^VP64^dCas9^VP64^) significantly increases activation of endogenous genes compared to a single copy of VP64 across various loci and cell types (Black et al., 2016, Chakraborty et al., 2014). In both screens, the cells were transduced with the gRNA library at a multiplicity of infection (MOI) of 0.2 to ensure one gRNA per cell, cultured for nine days and sorted via FACS for the 10% highest and lowest GFP-expressing cells. Deep sequencing of gRNA abundance in each population followed by differential expression analysis was used to identify enriched or depleted gRNAs in GFP-high or GFP-low cells.

### Modulation of paternal *SNRPN-2A-GFP* expression by dCas9^KRAB^

For the CRISPRi screen, the majority of significantly enriched or depleted gRNAs (gRNA “hits”) targeted the ∼300 kb high-density region (Figure 1D, Supp. Figure S1C). The CRISPRi screen of the paternal allele identified sites upstream, within, and downstream of the PWS-IC, and throughout the gene body of *SNRPN* (labelled pat1-pat8). Interestingly, the hits within and downstream of the *SNRPN* gene body had a strong DNA strand bias, with most hits targeted to the sense strand of the *SNRPN* gene (Supp. Figure S1D), potentially indicating steric hindrance of gene transcription (David et al., 2022).

When tested individually, gRNA hits from the paternal dCas9^KRAB^ screen all significantly influenced GFP expression, as assessed by mean fluorescence intensity (MFI) of the GFP reporter (Fig. 1F). Consistent with the expected repressive effect of dCas9^KRAB^, most gRNA hits in the paternal screen were enriched in the GFP-low population. However, several gRNAs within two regions, pat6 and pat8, increased GFP intensity (Figure 1F, Supp. Figure S1E). These regions are both downstream of *SNRPN* exon 10, with pat6 directly adjacent to exon 10 and pat8 ∼10 kb further downstream (Supp. Figure S1E). Surprisingly, targeting either of the two regions with dCas9^KRAB^ significantly raised GFP intensity but did not cause notable changes in the corresponding RNA expression measured by qRT-PCR (Supp. Figure S2A).

A previous study discovered that a weak polyadenylation (poly(A)) cleavage signal at the 3’ end of *SNRPN* enables continued transcription of downstream ncRNAs, including *SPA* lncRNAs and snoRNAs (Wu et al., 2016). We hypothesized that targeting dCas9^KRAB^ to the 3’ end of the *SNRPN* transcript might affect polyadenylation of the host transcript, thus leading to changes in SNRPN protein expression without altering transcription. In support of this hypothesis, targeting dCas9^KRAB^ to pat6 or pat8 regions increased polyadenylated *SNRPN* (Supp. Figure S2B) without altering total *SNRPN* RNA and down-regulated certain ncRNAs downstream of *SNRPN*, including *SPA1*, *SPA2* and *SNORD116* (Supp. Figure S2C). This represents a unique application of CRISPR-based screens to uncover possible alternative transcriptional termination sites. To assess potential changes to the 3’ untranslated region (UTR) of the *SNRPN* transcript, we performed 3’ RACE-sequencing (Supp. Figure S2D). However, there was a strong correlation in the abundance of SNRPN 3’ UTR sequence variants across all conditions, including dCas9^KRAB^ targeting and control groups, indicating no change in the SNRPN 3’ UTR (Supp. Figures S2E-G). This indicates that targeting dCas9^KRAB^ to pat6 and pat8 increased transcriptional termination and polyadenylation using the naturally functioning termination sequences.

### Activation of maternal *SNRPN-GFP* with ^VP64^dCas9^VP64^

Fewer hits were identified with the CRISPRa screen of the maternal allele. Two distinct clusters of gRNAs (labelled mat1 and mat2) were identified ∼100 kb upstream of the PWS-IC in the general region of annotated upstream SNRPN exons (Figure 1D). Interestingly, there was no overlap in the gRNA hits between the maternal and paternal screens (Supp. Figure S2H).

For CRISPRa validations, we first used pools of three or four gRNAs targeting the same region to leverage the synergistic activity of multiple gRNAs (Maeder et al., 2013, Perez-Pinera et al., 2013). Targeting both mat1 and mat2 regions with ^VP64^dCas9^VP64^ significantly up-regulated *matSNRPN-2A-GFP*, as assessed by qRT-PCR (Figure 1G). Individual validation of gRNAs within each pool revealed that the single best gRNAs for each region were mat1 g3 and mat2 g2, which we used for subsequent studies with ^VP64^dCas9^VP64^ (Figure 1G). Interestingly, pooling gRNAs in the mat1 region improved *SNRPN-GFP* activation, while pooling gRNAs in the mat2 region resulted in weaker activation compared to using mat2 g2 alone. This may be explained by the overlap of genomic locations among the three mat2 gRNAs, which could hinder dCas9 binding when multiple gRNAs are present in the same cell. This was not the case with the four mat1 gRNAs (Supplementary Table 1).

To ensure that the lack of overlap between the maternal and paternal screens was not due to allele-specific sensitivity bias in GFP reporters, we tested gRNA hits individually in cells with the opposite reporter allele. Most gRNAs showed no influence on *SNRPN* expression on the other allele (Supp. Figures S2I-J). Region pat5, overlapping the PWS-IC, had a minor effect (∼20-fold) on maternal *SNRPN* expression when perturbed with ^VP64^dCas9^VP64^ (Supp. Figure S2J), which was modest compared to fold-changes observed at mat1 and mat2 regions (Figure 1G).

### Activation of maternal *SNRPN-2A-GFP* via targeted demethylation

We hypothesized that the lack of overlap between hits in the CRISPRa and CRISPRi screens of the maternal and paternal alleles was due in part to differences in the epigenetic mechanisms regulating the identified elements. VP64 activates gene expression through the recruitment of epigenetic modifiers and transcriptional machinery, often establishing a more accessible chromatin environment enriched in active histone marks, such as H3K27ac and H3K4me3 (Gao et al., 2014, Gao et al., 2013, Hilton et al., 2015). However, VP64 may not fully investigate the influence of other epigenetic modifications, such as DNA methylation, linked to imprinting at the 15q11-13 locus. To directly assess the role of DNA methylation in maintaining the maternal imprint, we used dCas9 fused to the catalytic domain of Ten-eleven translocation methylcytosine dioxygenase 1 (^Tet1c^dCas9) to catalyze DNA demethylation at the target locus. This fusion can demethylate DNA in a targeted manner and induce corresponding changes in gene expression (Josipović et al., 2019, Liu et al., 2016).

We performed a similar screen with ^Tet1c^dCas9 as we had completed with ^VP64^dCas9^VP64^, sorting cells based on *matSNRPN-2A-GFP* expression (Figure 1C,D). In this case, we designed a gRNA sub-library comprising all the hits from the previous three screens, totaling 583 gRNAs, including 50 non-targeting controls (Figure 1E). The sub-library was designed to enable small-scale screens to assess the impact of various dCas9-based epigenome editing effectors, including ^Tet1c^dCas9, on different chromatin marks at the regulatory regions identified in the initial CRISPRa and CRISPRi screens. The sub-library screen with ^VP64^dCas9^VP64^ confirmed hits within the mat1 and mat2 regions, albeit with higher sensitivity (Figure S1C, S2K), providing validation of and adding confidence to the screening methods. The screen with ^Tet1c^dCas9 identified gRNA hits in the PWS-IC, overlapping a CpG island within *SNRPN* exon 1 (mat3), which were not detected with ^VP64^dCas9^VP64^ (Figure 1H). These gRNA hits largely overlapped with significant hits in the pat4 region from the dCas9^KRAB^ screen (Figure 1D, Supp. Figure S2L). We validated 8 individual gRNA hits in the PWS-IC, which showed robust activation of mat*SNRPN-GFP* (Figure 1I).

### Activation of maternal PWS genes in isogenic wildtype and PWS iPSC lines

To assess how ^Tet1c^dCas9-or ^VP64^dCas9^VP64^-mediated maternal PWS gene activation compares to PWS gene expression from the paternal allele in the absence of any reporter construct, we tested several effector-gRNA combinations in isogenic wildtype (WT) iPSCs and iPSCs with a PWS Type II deletion (ΔPWS) introduced via Cas9 nuclease (Figure 2A). Stable ^VP64^dCas9^VP64^ and ^Tet1c^dCas9 cell lines were generated and transduced with individual gRNAs chosen from screen validations, and transcriptome-wide gene expression was assessed at 14 days post-transduction by total RNA-seq, as well as RT-qPCR for *SNRPN* and *SNORD116*.

**Figure 2.**
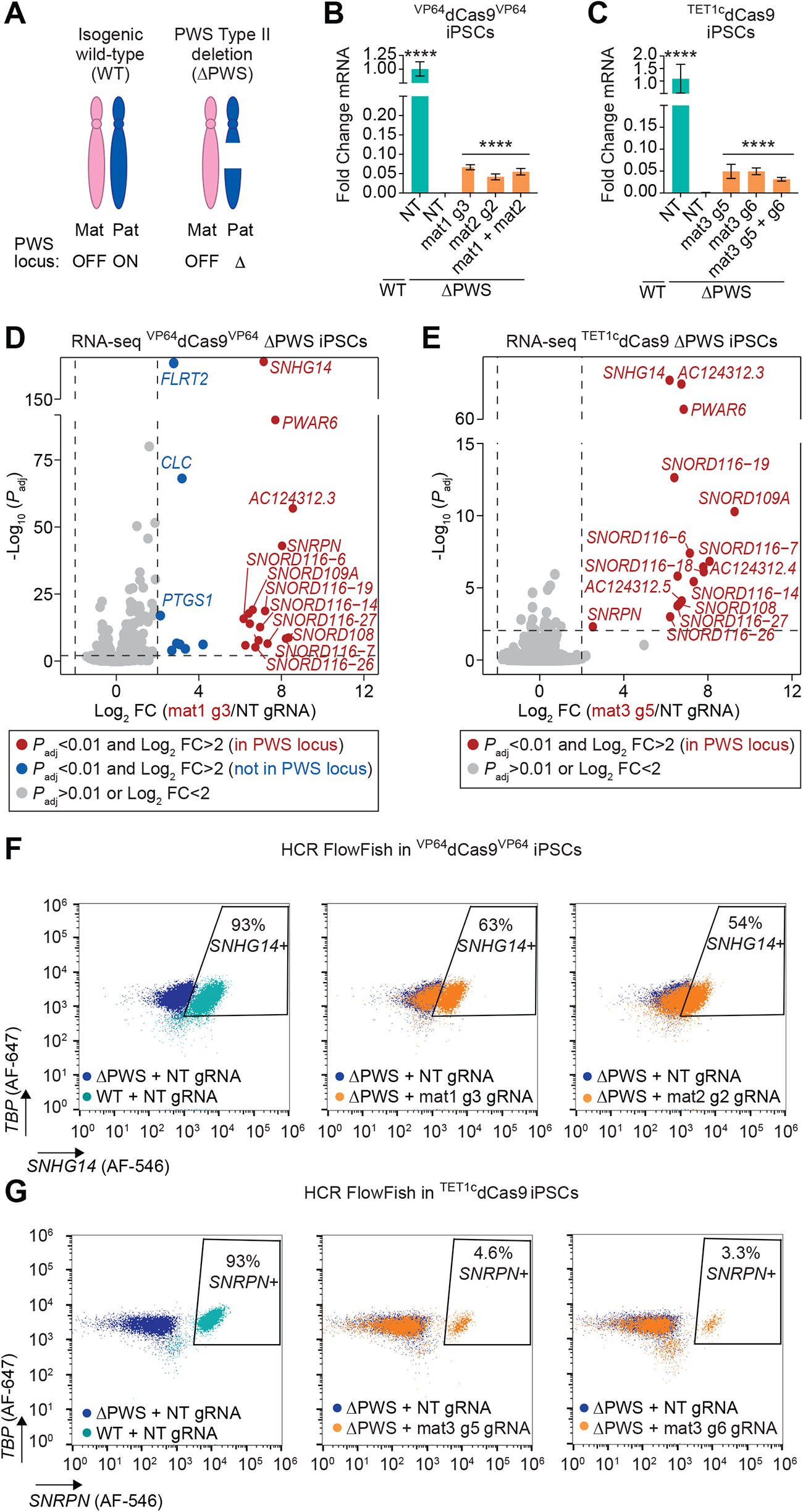
Tet1c and VP64 activate maternally imprinted PWS genes in ΔPWS iPSCs. (A) Schematic of chr15 in isogenic wildtype (WT) and PWS Type II deletion (ΔPWS) iPSCs. (B) qPCR of *SNRPN* in WT or ΔPWS iPSCs with ^VP64^dCas9^VP64^ 14 days after transduction with the indicated gRNA or gRNA pool. (C) qPCR of *SNRPN* in WT or ΔPWS iPSCs with ^Tet1c^dCas9 14 days after transduction with the indicated gRNA or gRNA pool. For both qPCR plots, fold change values are plotted mean +/-SD, but statistics were calculated on ddCt values (normalized to *GAPDH* and WT ctrl sample); one-way ANOVA followed by Dunnett’s test vs. ΔPWS NT gRNA ****p<0.0001. (D) Differential expression analysis of total RNA sequencing of ^VP64^dCas9^VP64^ ΔPWS iPSCs, comparing mat1 g3 to NT gRNA (E) Differential expression analysis of total RNA sequencing of ^Tet1c^dCas9 ΔPWS iPSCs, comparing mat3 g5 to NT gRNA. (F) HCR FlowFISH of ^VP64^dCas9^VP64^ iPSCs (WT or ΔPWS) with the indicated gRNA. *SNHG14* signal on X axis, with *TBP* as a control for cell size and staining. (G) HCR FlowFISH of ^Tet1c^dCas9 iPSCs (WT or ΔPWS) with the indicated gRNA. *SNRPN* (transcript variant 1) signal on X axis, with *TBP* as a control for cell size and staining.

Both ^VP64^dCas9^VP64^ and ^Tet1c^dCas9, in combination with a single gRNA, specifically activated PWS transcripts downstream of the gRNA binding sites, including *SNRPN*, *snoRNA116* (*SNORD116*) transcripts, and the *SNRPN* and *SNORD116/115* long host transcripts (*AC124312* and *SNHG14*), with minimal off-target effects (Figure 2B-E). In ^VP64^dCas9^VP64^ ΔPWS iPSCs, 8 genes outside the PWS locus showed significant differential expression, although the changes were lower than any PWS genes (Figure 2D). It is possible that these expression changes outside the PWS locus are consequences of upregulation of PWS genes, as several PWS lncRNAs, including *IPW* in particular, are known to regulate the expression of other genes in trans (Deshpande et al., 2022, Stelzer et al., 2014, Sledziowska et al., 2023). However, we were unable to find any reported evidence that *IPW* or other PWS lncRNAs regulate these differentially expressed genes. While both ^VP64^dCas9^VP64^ and ^Tet1c^dCas9 robustly activated maternal *SNRPN*, expression only reached approximately 5-10% of wildtype levels as assessed by RT-qPCR (Figure 2B,C). This indicates either incomplete reactivation of *SNRPN* or complete reactivation in only a subset of cells. Similarly, ^VP64^dCas9^VP64^ and ^Tet1c^dCas9 activated downstream transcript *SNORD116* to about 10-30% of wildtype expression, with ^VP64^dCas9^VP64^ having a stronger effect (Supp. Figure S3A,B). Interestingly, ^VP64^dCas9^VP64^ with mat1 g3 induced transcription upstream of the PWS-IC but not at the canonical exon 1 of *SNRPN* in both WT and ΔPWS iPSCs, whereas ^Tet1c^dCas9 with mat3 g5 induced transcription at *SNRPN* exon 1, as assessed by RT-qPCR of exon junctions specific to different subsets of transcripts (Supp. Figure S3C,D).

RNA sequencing further supported ^VP64^dCas9^VP64^ activation of transcript variants initiating upstream of the PWS-IC, in the exon immediately downstream of the mat1 g3 gRNA binding site (Supp. Figure S4A). Notably, these *SNRPN* transcripts initiating at upstream exons are normally expressed in neurons but not in most other somatic cell types (Runte et al., 2001, Landers et al., 2004), suggesting that ^VP64^dCas9^VP64^ with mat1 g3 might recapitulate cell type-specific regulation. Additionally, ^VP64^dCas9^VP64^ at the mat1 region decreased transcripts containing exon 1 in WT cells (Supp. Figure S3C), suggesting that the transcription starting at upstream exons and proceeding through the canonical promoter region on the paternal allele may disrupt normal *SNRPN* transcription.

To assess the proportion of PWS cells expressing maternal *SNRPN*, we used HCR-FlowFish (Reilly et al., 2021) to stain for *SNHG14* transcript induced by ^VP64^dCas9^VP64^ and *SNRPN* transcript variant 1 induced by ^Tet1c^Cas9. These two different transcripts were measured because ^VP64^dCas9^VP64^ and ^Tet1c^dCas9 preferentially upregulate different transcript variants (Supp. Figure S3A-D). Cells were sequentially transduced with lentiviral vectors encoding dCas9-effector and gRNA expression cassettes, followed by antibiotic selection. ^VP64^dCas9^VP64^ with either mat1 g3 or mat2 g2 gRNAs modestly increased *SNHG14* transcript in most cells, whereas ^Tet1c^dCas9 with gRNAs mat3 g5 or mat3 g6 fully restored *SNRPN* transcripts to wildtype levels but only in only 5% (g5) or 3% (g6) of cells (Figure 2F,G; Supp. Figure S3E,F). This finding was further corroborated by DNA methylation analysis via bisulphite sequencing, which showed a reduction in DNA methylation levels at a CpG-rich region within the PWS-IC with ^Tet1c^dCas9 but not ^VP64^dCas9^VP64^ (Figure 3A,B, Supp. Figure S4B). Interestingly, while there was a slight reduction in DNA methylation in this region with ^Tet1c^dCas9 at 15 days post-transduction (Supp. Figure S4B), the demethylation was more pronounced at a later timepoint, 38 days post-transduction (Figure 3B). At this later timepoint, *SNRPN* expression levels were approximately 5% of wildtype with ^Tet1c^dCas9 and mat3 g5 (Supp. Figure S4C), corresponding to the observed ∼5% demethylation of the maternal PWS-IC. Taken together, these results indicate that demethylation of the PWS-IC on the silenced maternal locus can completely restore *SNRPN* transcription.

**Figure 3.**
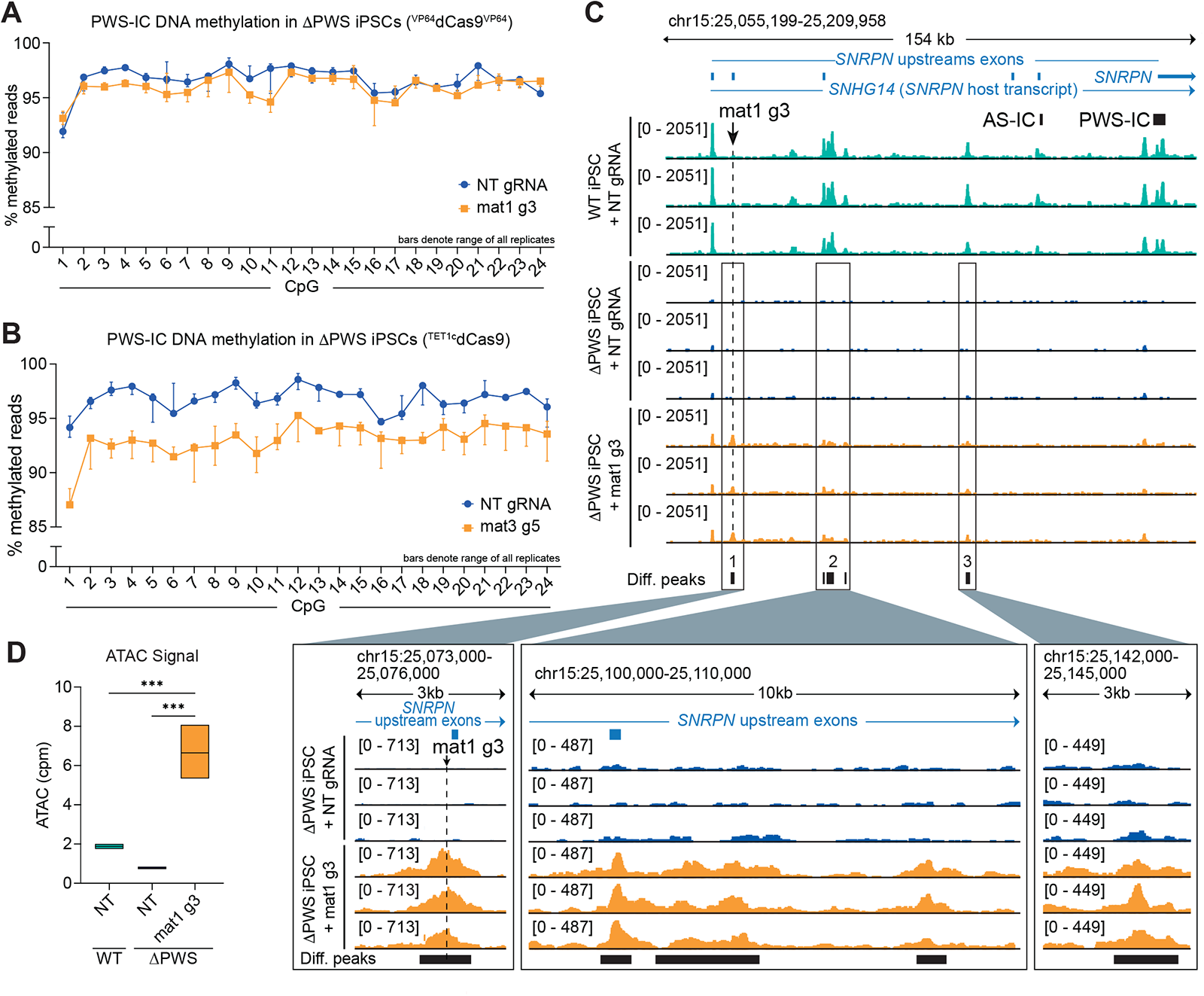
Tet1c and VP64 alter chromatin accessibility and/or DNA methylation at the PWS locus. (A) Targeted bisulphite sequencing of ΔPWS iPSCs with ^VP64^dCas9^VP64^ covering 24 CpG sites within the PWS locus (hg19 chr15: 25200353-25200693), 38 days post-transduction. (B) Targeted bisulphite sequencing of ΔPWS iPSCs with ^Tet1c^dCas9 covering 24 CpG sites within the PWS locus (hg19 chr15: 25200353-25200693), 38 days post-transduction. Data for (A) and (B) are shown as the range of the data, with the plotted point being the median, n = 3 replicates. (C) Browser tracks of ATAC sequencing (rpkm-normalized BigWig) of ΔPWS or WT iPSCs with ^VP64^dCas9^VP64^ and NT or mat1 g3 gRNA. (D) Quantification of ATAC-seq reads (counts per million) at the peak at the mat1 g3 guide binding site (dashed line in (A)). ***p < 0.001, Tukey’s test following one-way ANOVA.

We used ATAC-sequencing to assess changes in chromatin accessibility in ΔPWS iPSCs following treatment with ^VP64^dCas9^VP64^ or ^Tet1c^dCas9 (Corces et al., 2017). With ^VP64^dCas9^VP64^ and mat1 g3 gRNA, we observed six differential peaks (p_adj_ < 0.01 and log_2_(fold-change) > 1) in ΔPWS iPSCs compared to treatment with a control non-targeting gRNA, all within the PWS locus. Five of these peaks were located upstream of *SNRPN,* within the *SNRPN* host transcript *SNHG14* (Figure 3C), and the sixth peak was in the *PWAR1* gene, approximately 160 kb downstream of *SNRPN*, within the *SNHG14* host transcript that transcribes through the *SNORD116/115* cluster (Supp. Figure S3G). ^VP64^dCas9^VP64^ increased chromatin accessibility at the gRNA binding site, creating an ATAC peak not detected in WT cells (Figure 3D). Downstream of the gRNA binding site, the chromatin accessibility landscape in ΔPWS iPSCs with ^VP64^dCas9^VP64^ and mat1 g3 gRNA resembled that of WT iPSCs, albeit with smaller peaks, likely due to the heterogeneity in the extent of VP64-induced chromatin remodeling and/or *SNRPN* activation in the cell population, as indicated by the HCR-FlowFISH results (Figure 2F). Collectively, these unbiased and comprehensive profiling assays demonstrate that ^VP64^dCas9^VP64^ specifically facilitates transcription and increases chromatin accessibility throughout the PWS locus in a manner that recapitulates the paternal locus of WT cells, with minimal off-target effects.

In contrast, there were no differential chromatin accessibility peaks detected in ΔPWS iPSCs with ^Tet1c^dCas9 and mat3 g5 gRNA compared to non-targeting gRNA (Supp. Figure S4D), although a slight increase in ATAC signal was observed at the PWS-IC (Supp. Figure S4E). This might be due to the assay’s limited sensitivity to detect changes in chromatin accessibility in these samples, as the FlowFISH and bisulphite sequencing data showed that ^Tet1c^dCas9 demethylated the PWS-IC and activated *SNRPN* expression in only 5% of the cells. Nonetheless, these results support a model in which targeted activation with VP64 and demethylation by Tet1c modulate the locus through distinct epigenetic mechanisms.

### Transient delivery of ^TET1c^dCas9 to PWS iPSCs induces stable, heritable activation of PWS genes

Deposition of DNA methylation at a promoter by dCas9-DNMT3 can induce stable silencing of target genes either alone or in combination with KRAB (Bintu et al., 2016, Nuñez et al., 2021, Amabile et al., 2016). Conversely, DNA demethylation mediated by ^TET1^dCas9 can activate genes with methylation-sensitive promoters (Liu et al., 2016, Nuñez et al., 2021). Because the transcriptional machinery that normally maintains expression of PWS genes from the paternal allele should be present, we postulated that demethylation of the maternal PWS-IC at an initial time point by transient expression of ^TET1^dCas9 might be sufficient to stably activate the silenced PWS locus.

In an attempt to improve the efficiency of TET1-mediated *SNRPN* activation, we compared three different TET1 constructs via lentiviral transduction: the original ^TET1c^dCas9 construct, SunTag TET1c (modified from (Morita et al., 2016)), and TET1v4 dCas9 (modified from (Nuñez et al., 2021)). SunTag TET1c uses a recruitment strategy to recruit up to 5 copies of TET1c to a single Cas9 (Morita et al., 2016). TET1v4, like the original ^TET1c^dCas9 construct, is a direct fusion of TET1c to the N-terminus of dCas9; however, TET1v4 uses a longer 80-amino acid XTEN80 linker (Nuñez et al., 2021), compared to the original 49-amino acid linker. The latter two TET1 systems were cloned into a lentiviral backbone and modified with selectable markers for stable transduction. Among the tested TET1 systems, TET1v4 showed the strongest activation of *SNRPN* in ΔPWS iPSCs (Supp. Figure S4F). We further engineered TET1v4 into a smaller plasmid for efficient transient transfection and replaced the selectable marker with Thy1.1 (CD90), a surface protein that enables robust and sensitive antibody staining to sort cells receiving the transgene.

We used nucleofection to transiently deliver the TET1v4-dCas9-T2A-Thy1.1 (^TET1v4^dCas9) plasmid to WT and ΔPWS iPSCs stably expressing the gRNA (delivered via lentivirus) (Figure 4A). We sorted nucleofected cells at 2 days post-nucleofection on Thy1.1 reporter expression, ensuring that the cells we assayed received both ^TET1v4^dCas9 and gRNA. With this method, dCas9 expression in the cells was undetectable by day 8 post-nucleofection (Figure 4B). However, *SNRPN* expression in the ΔPWS iPSCs nucleofected with ^TET1v4^dCas9 plasmid increased and remained stable through 7 weeks post-nucleofection, indicating that transient expression of ^TET1v4^dCas9 is sufficient to stably and heritably reverse the silenced status of the PWS locus.

**Figure 4.**
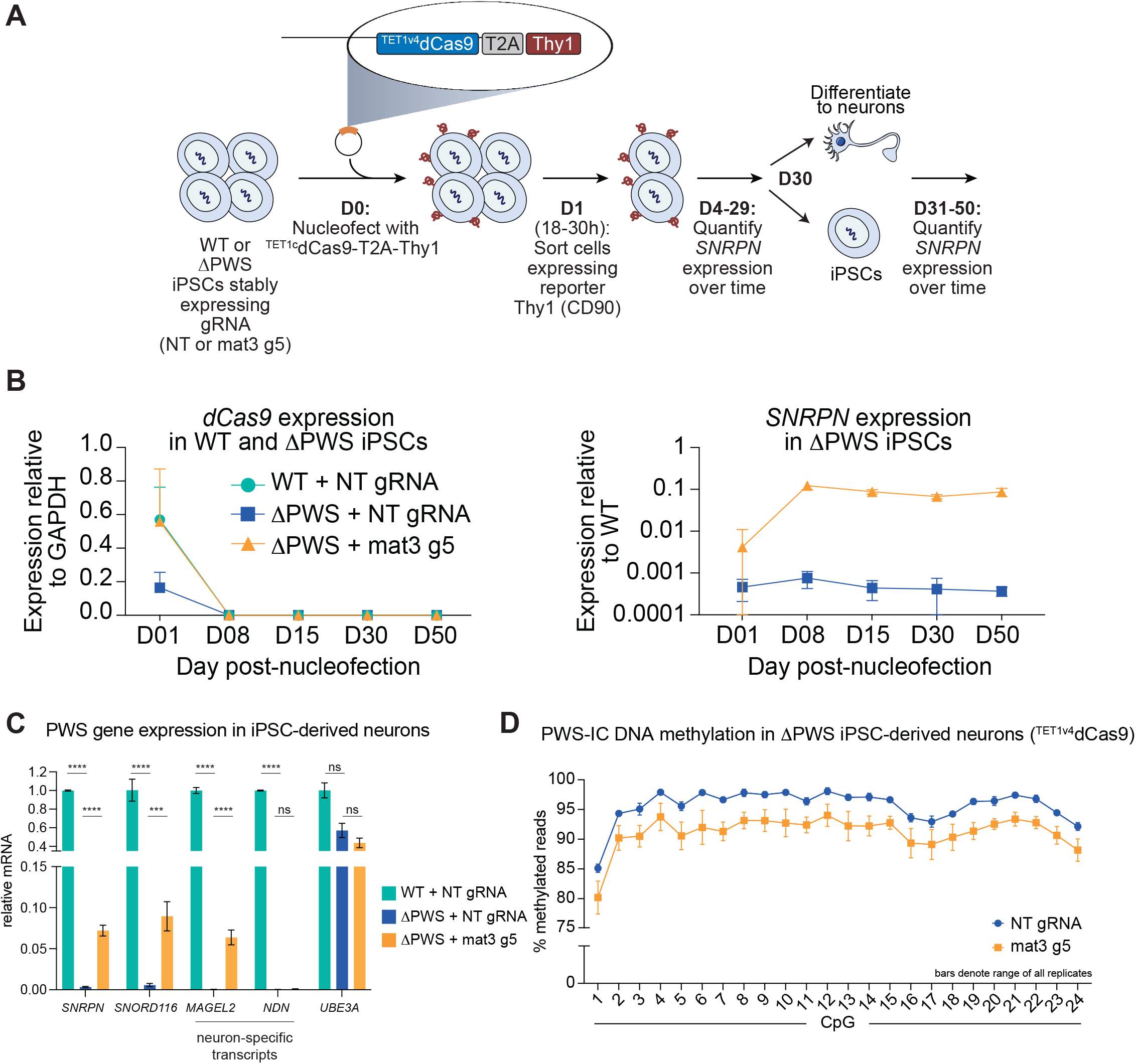
Transient expression of ^Tet1v4^dCas9 in ΔPWS iPSCs stably activates maternal PWS genes. (A) Schematic of experimental protocol for transient delivery of ^Tet1v4^dCas9 plasmid and PWS gene expression analysis. (B) qPCR of *dCas9* or *SNRPN* in WT or ΔPWS iPSCs after transient delivery of ^Tet1v4^dCas9 on Day 0. (C) qPCR of PWS genes in iPSC-derived neurons. Data plotted as mean fold change +/-SD, but statistics computed on ddCt (normalized to GAPDH and WT + NT). Two-way ANOVA followed by Dunnett’s test, compared to ΔPWS + NT gRNA; **p<0.01, ***p<0.001, ****p<0.0001. (D) Targeted bisulphite sequencing of ΔPWS iPSC-derived neurons approximately 21 days post-differentiation, covering 24 CpG sites within the PWS locus (hg19 chr15: 25200353-25200693). Data shown as median +/-range, n = 3 replicates.

We investigated the stability of the reactivated PWS locus during neuronal differentiation by differentiating ΔPWS iPSCs into neurons via Ngn2 overexpression at day 30 post-nucleofection (Figure 4A). At three weeks post-differentiation, *SNRPN* expression remained stably activated after differentiation into neurons (Figure 4C). Notably, maternal *SNRPN* expression in the ΔPWS iPSC-derived neurons showed comparable levels to those observed in the undifferentiated ΔPWS iPSCs transduced with ^TET1c^dCas9 and mat3 g5 (Figure 4C and Figure S3D), indicating no loss of *SNRPN* expression through differentiation or prolonged culture. Additionally, we examined the expression of neuron-specific imprinted PWS genes in these cells. Remarkably, the ΔPWS iPSC-derived neurons that received ^TET1v4^dCas9 and mat3 g5 showed a significant increase in the expression of *MAGEL2*, a transcript located 1.3 Mbp upstream of the PWS IC, to approximately 10% of wildtype levels (Figure 4C). Another neuron-specific imprinted PWS gene, *NDN*, located near *MAGEL2*, displayed an approximately two-fold upregulation, albeit still well below wildtype expression levels (Figure 4C).

Transcription of *SNRPN* host transcript *SNHG14* (also known as *UBE3A* antisense transcript) is known to suppress *UBE3A* (Hsiao et al., 2019, Wolter et al., 2020), which is located approximately 350kb downstream of *SNRPN. UBE3A* is biallelically expressed in most tissues but maternally expressed and paternally imprinted in neurons (Hsiao et al., 2019), and loss of *UBE3A* expression causes Angelman Syndrome (Clayton-Smith and Laan, 2003). We suspected that activating *SNRPN* from the maternal allele in neurons might impede normal transcription of *UBE3A.* Notably, *UBE3A* expression was not significantly decreased in iPSC-derived neurons that received ^TET1v4^dCas9 and mat3 g5 compared to cells that received ^TET1v4^dCas9 and NT gRNA (Figure 4C). However, *UBE3A* expression in the ΔPWS neurons, which retain only the maternal copy, was approximately halved compared to its expression in the WT neurons, which have both copies (Figure 4C), suggesting that *UBE3A* is incompletely imprinted in these cells. Incomplete *Ube3a* imprinting, resulting in paternal *Ube3a* expression, is observed in immature murine neurons (Judson et al., 2014, Jones et al., 2016). Given that iPSC-derived neurons differentiated via *Ngn2* overexpression resemble immature cortical neurons (Zhang et al., 2013), the apparent biallelic *UBE3A* expression we observed may be indicative of an immature molecular phenotype in these cells. Nonetheless, activation of PWS genes via demethylation of the PWS-IC on the maternal allele does not appear to disrupt *UBE3A* expression in ΔPWS iPSC-derived neurons.

Moreover, we analyzed DNA methylation at the PWS-IC in these ΔPWS iPSC-derived neurons (Figure 4D). DNA methylation at a region within the PWS-IC was approximately 5% lower in the ΔPWS iPSC-derived neurons that initially received ^TET1v4^dCas9 and mat3 g5 compared to non-targeting gRNA (Figure 4D), suggesting that stable demethylation was maintained through differentiation. These findings collectively indicate that the transient expression of ^TET1v4^dCas9 during early stages of cellular development is sufficient to establish stable and sustained reactivation of the silenced PWS locus, even through the process of neuronal differentiation.

## Discussion

In this study, CRISPR-based epigenetic screens at the 15q11-13 locus identify regulatory elements controlling expression of the *SNRPN* host transcript implicated in PWS. Targeting these regulatory elements with dCas9-based epigenetic editors leads to robust changes in gene expression of several candidate PWS-associated genes. This work provides proof-of-concept for a therapeutic strategy for PWS by reactivating maternal gene expression at the 15q11-13 locus through targeted dCas9-based epigenetic editing.

Because the PWS locus is imprinted, we generated allele-specific *SNRPN* reporter lines to enable independent analysis of maternal or paternal *SNRPN* gene expression. dCas9^KRAB^ repressed paternal *SNRPN* when targeted throughout the promoter and gene body, except at two regions at the 3’ end of the gene that unexpectedly increased overall *SNRPN* expression. Consistent with the hypothesis that dCas9^KRAB^ at the 3’ end of the gene alters its polyadenylation, we observed an increase in polyadenylated *SNRPN* as well as a decrease in downstream transcripts *SPA1*, *SPA2*, and *SNORD116*. Interestingly, we found that ^VP64^dCas9^VP64^ less effectively activated maternal *SNRPN* when targeted to the canonical promoter, the PWS-IC. DNA methylation may impede activation (Amabile et al., 2016, Devesa-Guerra et al., 2020), which could limit the effectiveness of VP64 at the methylated *SNRPN* promoter. However, we identified two regions located within *SNRPN* upstream exons that enabled upregulation of alternative *SNRPN* transcripts with ^VP64^dCas9^VP64^. Targeting ^VP64^dCas9^VP64^ to the mat1 region upstream of the *SNRPN* promoter effectively activated maternal *SNRPN* in a DNA-methylation independent manner. In contrast, ^TET1c^dCas9-mediated targeted DNA demethylation of the PWS-IC successfully activated maternal *SNRPN.* Different epigenome or transcriptional modifiers may function at different regulatory elements, depending on the underlying epigenetic landscape. Initiating transcription of upstream *SNRPN* exons may be important for brain-specific effects, as suggested by a patient presenting with PWS-like symptoms, with an unusual small deletion within the *SNRPN* upstream exons (Koufaris et al., 2016). Furthermore, in our study, ^VP64^dCas9^VP64^ exhibited stronger upregulation of *SNORD116* compared to ^TET1c^dCas9, possibly due to its location within the *SNRPN* upstream exons. This may reflect more effective upregulation of transcripts such as *SNHG14*, which extends through *SNRPN* and into downstream PWS genes.

Our findings reveal that transient delivery of ^TET1c^Cas9 to ΔPWS iPSCs results in stable activation of *SNRPN* that persists through neuronal differentiation. Furthermore, the imprinted gene *MAGEL2* is expressed in these neurons post-differentiation, indicating that DNA methylation is necessary for maintenance of the maternal PWS imprint. The PWS-IC is known to regulate expression of upstream neuronal transcripts (Chung et al., 2020). While previous studies have demonstrated stable gene repression induced by DNA methylation (Smith et al., 2008, Amabile et al., 2016, Nuñez et al., 2021), stable activation of multiple genes at an endogenous imprinted locus by removal of DNA methylation via transient delivery of a targeted DNA demethylase in order to compensate for disease-causing deletions is a unique application of these tools.

Unexpectedly, although we initially sorted cells that expressed both the gRNA and ^TET1c^Cas9 cassettes after nucleofection, only a portion of that cell population expressed mat*SNRPN*. Similarly, stable lentiviral delivery of the TET1 construct and gRNA, followed by drug selection for cells expressing both cassettes, also led to only a subset of the cells displaying mat*SNRPN* expression. There are several possible explanations for these observations. Firstly, we suspect that ^TET1c^Cas9 and/or gRNA expression levels might determine the efficiency of activation of the target gene, with a minimum expression threshold required to successfully demethylate the PWS-IC. Therefore, variation in expression levels among cells may account for the stochastic locus activation. Alternatively, as iPSC populations comprise a heterogeneous cell population (Narsinh et al., 2011), it is possible that TET1-mediated DNA demethylation is sufficient to induce *SNRPN* expression in a particular subset of cells, while others might require perturbation of additional chromatin marks and/or transcription factors to create a more transcriptionally permissive environment.

Our work builds upon other studies that have used targeted DNA demethylation editing to restore silenced gene expression in stem cells and neurons, including *BDNF*, *FMR1*, *MECP2*, and the imprinted *Dlk1-Dio3* locus (Liu et al., 2016, Liu et al., 2018, Qian et al., 2023, Kojima et al., 2022). These studies have demonstrated that targeting a dCas9-TET1 fusion to the methylated promoter of the silenced gene is sufficient to activate its expression. A recent report of activation of the MECP2 locus indicated that while targeted demethylation was sufficient for gene activation in iPSCs, additional changes to chromatin structure were necessary to achieve similar effects in neurons (Qian et al., 2023). It is possible that similar, further chromatin modifications are required to activate the silenced PWS locus in neurons compared to stem cells.

Our discovery that transient expression of ^TET1c^dCas9 stably and heritably activates PWS gene expression establishes the possibility of a one-time treatment for PWS. However, delivering large dCas9-based effectors with non-viral vectors remains a significant challenge *in vivo*. Additionally, because PWS is a complex disorder that affects the pituitary-hypothalamic axis and manifests with several co-morbidities including sleep disturbances (Irizarry et al., 2016), improvement of symptoms may require targeting multiple cell and tissue types *in vivo*. Thus, translating epigenome editing therapies for PWS to the clinic will necessitate further development of safe and efficient delivery of dCas9-based epigenome editors.

Further studies investigating the phenotypic effects of restoring PWS gene expression in animal models or patient-derived organoids would be valuable in establishing the potential of DNA methylation editing as a treatment for PWS. For instance, PWS patient-derived hypothalamic organoids have been shown to exhibit impaired leptin response (Huang et al., 2021), making them an ideal model for assessing the impact of dCas9-based epigenome editing to restore PWS gene expression in a cell type that is more disease-relevant. Future efforts will focus on directly targeting epigenome editing in neurons to restore PWS gene expression in order to better understand the relationship of PWS gene expression and PWS-related neurological phenotypes.

## Acknowledgments

We thank Lisa Burnett, Yong-Hui Jiang, Keith Siklenka, and all members of the Gersbach lab, especially S. Alexandra Garcia-Moreno, for technical assistance and helpful discussions. This work was supported by the Foundation for Prader Willi Research, Levo Therapeutics, and National Institutes of Health grants R01DA036865, U01AI146356, UM1HG013053, RM1HG011123, R01NS093200, and R01HD096326, National Science Foundation grants EFMA-1830957, DARPA grant HR0011-19-2-0008, an Allen Distinguished Investigator Award from the Paul G. Allen Frontiers Group to C.A.G, and the Open Philanthropy Project. D.R. was supported by a National Science Foundation Graduate Research Fellowship. J.B.B. was supported by NIGMS Biotechnology Training Grant T32GM008555.

## Author Contributions

D.R., J.B.B., N.I., M.T., and C.A.G. designed experiments. D.R., J.B.B., S.R.M., D.J.M., D.J.C.T., C.E.E., X.N., and N.I. performed experiments. D.R., J.B.B., and A.B. analysed the data. D.R., J.B.B., and C.A.G. wrote the manuscript. All authors reviewed and revised the manuscript.

## Resource Availability

### Lead Contact

Further information and requests for resources and reagents should be directed to and will be fulfilled by the lead contact, Charles Gersbach.

### Materials Availability

All materials are available to the research community through Materials Transfer Agreements.

### Data and Code Availability

Data generated in high-throughput sequencing studies are available through NCBI GEO with accession no. GSE243185. Any additional information required to reanalyse the data reported in this paper is available from the lead contact upon request.

## Methods

### Generation of *SNRPN-2A-GFP* pluripotent stem cell lines

A human iPS cell line (RVR-iPSCs) was used to construct the maternal and paternal *SNRPN-2A-GFP* reporter lines. RVR-iPSCs were retrovirally reprogrammed from BJ fibroblasts and characterized previously (Lee et al., 2012) . To generate the *SNRPN-2A-GFP* reporter lines, 3 x 10^6^ cells were dissociated with Accutase (Stemcell Tech, 7920) and electroporated with 6 μg of gRNA-Cas9 expression vector and 3 μg of *SNRPN* targeting vector using the P3 Primary Cell 4D-Nucleofector Kit (Lonza, V4XP-3032). Transfected cells were plated into a 10 cm dish coated with Matrigel (Corning, 354230) in complete mTesR1 (Stemcell Tech, 85850) supplemented with 10 μM Rock Inhibitor (Y-27632, Stemcell Tech, 72304). 24 h after transfection, positive selection began with 1 μg/mL puromycin for 7 days. Following selection, cells were transfected with a CMV-CRE recombinase expression vector to remove the floxed puromycin selection cassette. Transfected cells were expanded and plated at low density for clonal isolation (180 cells/cm^2^). Resulting clones were mechanically picked and expanded, and gDNA was extracted using QuickExtract DNA Extraction Solution (Lucigen, QE09050) for PCR screening of targeting vector integration. A polyclonal cell line expressing lentivirally transduced dCas9^KRAB^, ^VP64^dCas9^VP64^ or ^Tet1c^dCas9 was used for the CRISPR screens and validations.

### Generation of Type II PWS iPSC lines

We used our previously developed Single-guide-CRISPR/Cas-targeting-Of-Repetitive-Elements (SCORE) method (Tai et al., 2016) to generate Type II PWS deletion in isogenic human iPSCs, MGH2069 cells (Seo et al., 2017). We designed guide RNAs (gRNAs) targeting specific pair of segmental duplication blocks (BP2 and BP3) at the PWS locus (target sequence 5′-CTCCCTGCCTAGAAGCTGGT-3′, chromosome 15: 23,198,458–23,198,477 and 28,490,195–28,490,214). The details of the characterization and maintenance of PWS iPSC lines are described in our previous work (Chen et al., 2020).

### Cell culture

Human iPSCs were maintained in mTeSR1 (StemCell Tech, 85850) or mTeSR Plus (StemCell Tech, 100-0276) on Matrigel-coated tissue culture plastic in a 37C incubator with 5% CO2. Prior to use, plates were coated with Matrigel (Corning, 354230) at a concentration of 1mg per 24mL DMEM/F12 (Gibco, 11320033) and incubated for at least 1 hour at 37C. Medium was supplemented with 10uM ROCK inhibitor (ROCKi) (Y-27632, Stemcell Tech, 72304) for 16-48 hours when cells were passaged with Accutase (StemCell Tech, 07920; Innovative Cell Technologies, AT104; Gibco A1110501) and otherwise omitted.

### iPSC nucleofection and cell sorting

For plasmid nucleofection, iPSCs were dissociated from the plate with Accutase diluted 1:1 in divalent cation-free PBS. We used the Lonza P3 Primary Cell Nucleofector Solution X Kit L (Lonza, VXP-3024) for all nucleofection experiments. Cells were counted, spun down for 10 minutes at 100xg, and resuspended in Lonza P3 Primary Cell Nucleofector Solution at approximately 8 million cells per 100uL reaction. 10ug of plasmid were used for each 100uL reaction. Cells were nucleofected with the Nucleofector X Unit (Lonza) using programme CB-150. Samples were immediately resuspended in pre-warmed mTeSR Plus media with 10uM ROCKi and transferred to 6-well plates (4-8 million cells per well). After 18-24 hours in the incubator, cells were washed with PBS, and adherent cells were dissociated again with Accutase. Samples were stained in PBS with eBioscience Fixable Viability Dye eFluor 780 (Invitrogen, 65-0865-18) and Anti-Rat CD90 Antibody FITC (referred to in the text as mouse Thy1) (StemCell, 60024Fl.1, Lot no. 1000072622). Stained, untransfected cells were used as a negative control for the anti-CD90 antibody. Live, CD90-positive cells were sorted with the Sony SH800Z cell sorter into tubes containing DMEM/F12, 10uM ROCKi, and 1% antibiotic-antimycotic (Gibco, 15240062). Approximately 20% of the sorted cells were pelleted and flash-frozen for later RNA extractions. Approximately 80% of the sorted cells were pelleted for 5 minutes at 300xg, re-suspended in mTeSR Plus, 10uM ROCKi, and 1% antibiotic-antimycotic, and plated in 24-well plates. One or two days later, media were changed to mTeSR Plus without antibiotic-antimycotic. ROCKi was omitted from the media once colonies were well-established, usually two to three days after initial plating. Media were changed daily or once every two days.

### Neuronal differentiation

WT and ΔPWS iPSCs were differentiated via Ngn2 over-expression as described in a published protocol, with modifications (Wang et al., 2017). Briefly, on day -4, iPSCs were transduced during plating with lentiviruses encoding *TetO*-*mNgn2* and *hUBC-M2rtTA* to deliver doxycycline-responsive mouse *Ngn2* cDNA. After transduction, cells were kept in mTeSR Plus with 10uM ROCKi for one day. On day -3, media were changed to pre-differentiation media with 2ug/mL doxycycline, as described in the protocol. Media were changed daily. On day 0, pre-differentiated cells were dissociated and re-plated in post-differentiation media with 2ug/mL doxycycline at approximately 100,000 cells per cm^2^. Partial media changes occurred every 7 days (no doxycycline).

### Lentiviral production and titration

HEK293T cells were acquired from the American Tissue Collection Center (ATCC) and purchased through the Duke University Cell Culture Facility. The cells were maintained in DMEM High Glucose supplemented with 10% FBS and 1% penicillin-streptomycin and cultured at 37°C with 5% CO2. For lentiviral production of the gRNA libraries, ^VP64^dCas9^VP64^ and dCas9^KRAB^, 4.5 x 10^6^ cells were transfected using the calcium phosphate precipitation method (Salmon and Trono, 2007) with 6 μg pMD2.G (Addgene #12259), 15 μg psPAX2 (Addgene #12260) and 20 μg of the transfer vector. The medium was exchanged 12-14 h after transfection, and the viral supernatant was harvested 24 and 48 h after this medium change. The viral supernatant was pooled and centrifuged at 600*g* for 10 min, passed through a PVDF 0.45 μm filter (Millipore, SLHV033RB), and concentrated to 50x in 1x PBS using Lenti-X Concentrator (Clontech, 631232) in accordance with the manufacturer’s protocol. To produce lentivirus for gRNA validations or dCas9 effectors, 1.2 x 10^6^ cells were transfected in 6-well plates using Lipofectamine 3000 (Invitrogen, L3000008) according to the manufacturers instructions with 200 ng pMD2.G (Addgene #12259), 600 ng psPAX2 (Addgene #12260) and 500 ng of the transfer vector. The medium was exchanged 6 h after transfection, and the viral supernatant was harvested 24 and 48 h after transfection. The viral supernatant was pooled and filtered through a 0.45um filter or centrifuged for 10 min at 600xg to remove cell debris, then concentrated to 50x in 1x PBS using Lenti-X Concentrator (Takara, 631232), in accordance with the manufacturer’s protocol. The titer of the lentiviral gRNA library pool was determined by transducing 3 x 10^4^ cells with serial dilutions of lentivirus and measuring the percent mCherry expression 4 d after transduction with a SH800 FACS Cell Sorter (Sony Biotechnology). All lentiviral titrations were performed in the cell lines used in the CRISPR screens.

### RNA isolation and quantitative RT-PCR

Total RNA was isolated using RNeasy Plus (Qiagen, 74136) and QIAshredder kits (Qiagen, 79656) for VP64 and KRAB gRNA validations in SNRPN-2A-GFP iPSCs. Total RNA was isolated using Norgen Total RNA Purification Plus Micro kit (Norgen, 48500) for other experiments, including RNA sequencing. Reverse transcription was carried out on 0.1-0.5 μg total RNA per sample in a 10 or 20 μl reaction using the SuperScript VILO Reverse Transcription Kit (Invitrogen, 11754). 10 ng of cDNA was used per PCR reaction with Perfecta SYBR Green Fastmix (Quanta BioSciences, 95072) using the CFX96 Real-Time PCR Detection System (Bio-Rad). All amplicon products were verified by melting curve analysis. All qRT-PCR results are presented as fold change in RNA normalized to *GAPDH* expression. To purify poly(A) RNA, total RNA was first isolated using RNeasy Plus (Qiagen) as described above. Poly(A) RNA was purified from 1 μg total RNA using RNA purification beads as part of the Truseq Stranded mRNA kit according to the manufacturer’s protocol (Illumina). 1 ng of poly(A)-enriched RNA was loaded into each qRT-PCR reaction.

### RNA sequencing and data analysis

For RNA sequencing, total RNA isolated with the Norgen RNA Purification Plus Micro kit (Norgen, 48500) was submitted to Genewiz (Azenta Life Sciences) for total RNA sequencing. Genewiz verified quality of samples and libraries and performed DNAse treatment as necessary. Sequencing reads were trimmed with Trimmomatic to remove adapters and filter on read quality. Reads were mapped to hg19 with STAR. DeepTools bamCoverage was used to generate RPKM-normalized bedgraph files, then bedGraphToBigWig was used to convert bedgraph to bigwig files for visualization in the genome browser. Read counts for each gene were created using the featureCounts function from the subread package (v1.4.6-p4). We used DESeq2 for differential expression analysis, where gene counts are fitted to negative binomial general linearized models (GLMs) and Wald statistics to determine significantly differentially expressed genes.

### Plasmid Construction

The lentiviral ^VP64^dCas9^VP64^ plasmid was generated by modifying Addgene #59791 to replace *GFP* with *BSD* blasticidin resistance. The ^Tet1c^dCas9 plasmid was generated by amplifying Tet1c from Addgene #108245, followed by Gibson assembly of Tet1c, dCas9, and BSD into a lentiviral expression backbone. The dCas9^KRAB^ plasmid is equivalent to Addgene #67620 but with BSD replacing GFP, and the LacZ gene is not present in the backbone. The ^Tet1c^Cas9-T2A-Thy1.1 plasmid for transfection was generated by replacing *BFP* in Addgene #167983 with mouse *Thy1.1* (*CD90*). The gRNA expression plasmid for the single gRNA screens was generated by modifying Addgene #83925 to contain an optimized gRNA scaffold (Chen et al., 2013), a puromycin resistance gene in place of Bsr and a mCherry transgene in place of GFP. For creating stable gRNA-expressing cell lines in WT and ΔPWS iPSCs for nucleofection of the ^Tet1c^Cas9 effector, we used a gRNA expression plasmid that was further modified by removing the mCherry transgene. The gRNA expression plasmid for the dual gRNA screen was generated by further modification of the single gRNA expression plasmid to contain an additional gRNA cassette expressing sg-mat1 under control of the mU6 Pol III promoter with a modified gRNA scaffold described previously (Adamson et al., 2016). Individual gRNAs were ordered as oligonucleotides (Integrated DNA Technologies) and cloned into the gRNA expression plasmids using BsmBI sites. The *SNRPN* targeting vector was cloned by inserting ∼700 bp homology arms (surrounding the *SNRPN* stop codon in exon 10), amplified by PCR from genomic DNA of RVR-iPS cells, flanking a P2A–GFP sequence with a LoxP-puromycin resistance cassette.

### gRNA library design and cloning

For the full gRNA library, the gt-scan algorithm (O’Brien and Bailey, 2014) was used to identify all possible gRNAs within the chromosome 15q11-13 region and rank the gRNAs by off-target alignments to the human genome. DNase I hypersensitivity sites (DHSs) for H1 human embryonic stem cells (H1 hESCs) were downloaded from ENCODE (http://www.encodeproject.org). The high-density gRNA region spanned chr15:25,064,194-25,368,441 (hg19) and consisted of the top 30% of all gRNAs within this region ranked by off-target score. Outside of the high-density region and within chr15:23,692,325-26,425,399 (hg19), all gRNAs within DHS coordinates in H1 hESCs were selected for inclusion in the library. In total, 11,751 gRNAs were designed, including 531 non-targeting control gRNAs that were selected from a previous genome-wide CRISPRi gRNA library (Horlbeck et al., 2016). For the gRNA sub-library used in the ^Tet1c^dCas9 screen, all hits (adjusted p-value<0.05) from the previous CRISPRa and CRISPRi screens were included, in addition to 50 non-targeting control gRNAs, for a total of 583 gRNAs. For both gRNA libraries, the oligonucleotide pool (Twist Bioscience) was PCR amplified and cloned using Gibson assembly into the single or dual gRNA expression plasmid. The gRNA libraries are available for download in Supplemental Materials.

### dCas9^KRAB^ and ^VP64^dCas9^VP64^ SNRPN-2a-GFP screens

Each screen was performed in triplicate with independent transductions. For each replicate, 24 x 10^6^ *SNRPN-2A-GFP* ^VP64^dCas9^VP64^ (maternal) or dCas9^KRAB^ (paternal) iPSCs were dissociated using Accutase (Stemcell Tech, 7920) and transduced in suspension across five matrigel-coated 15-cm dishes in mTesR (Stemcell Tech 85850) supplemented with 10 μM Rock Inhibitor (Y-27632, Stemcell Tech, 72304). Cells were transduced at a MOI of 0.2 to obtain one gRNA per cell and ∼430-fold coverage of the gRNA library. The medium was changed to fresh mTesR without Rock Inhibitor 18-20 h after transduction. Antibiotic selection was started 30 h after transduction by adding 1 μg/mL puromycin (Sigma, P8833) directly to the plates without changing the medium. The cells were fed daily and passaged as necessary maintaining library coverage until harvest. Cells were harvested for sorting 9 d after transduction of the gRNA library for all three screens. Cells were washed once with 1x PBS, dissociated using Accutase, filtered through a 30 μm CellTrics filter (Sysmex, 04-004-2326) and resuspended in FACS Buffer (0.5% BSA (Sigma, A7906), 2 mM EDTA (Sigma, E7889) in PBS). Before sorting, an aliquot of 4.8 x 10^6^ cells were taken to represent a bulk unsorted population. The highest and lowest 10% of cells were sorted based on GFP expression and 4.8 x 10^6^ cells were sorted into each bin. Sorting was done with a SH800 FACS Cell Sorter (Sony Biotechnology). After sorting, genomic DNA was harvested with the DNeasy Blood and Tissue Kit (Qiagen, 69506).

### ^Tet1c^dCas9 and ^VP64^dCas9^VP64^ SNRPN-2A-GFP sub-library screens

The screen was performed in triplicate with independent transductions. For each replicate, 1.7 x 10^6^ *matSNRPN-2A-GFP* ^Tet1c^dCas9 or ^VP64^dCas9^VP64^ iPSCs were dissociated using Accutase (Stemcell Tech, 7920) and transduced in suspension in a matrigel-coated 10-cm dish in mTesR (Stemcell Tech 85850) supplemented with 10 μM Rock Inhibitor (Y-27632, Stemcell Tech, 72304). Cells were transduced at a MOI of 0.2 to obtain one gRNA per cell and ∼580-fold coverage of the gRNA sub-library. The medium was changed to fresh mTesR without Rock Inhibitor 18-20 h after transduction. Antibiotic selection was started 30 h after transduction by adding 1 μg/mL puromycin (Sigma, P8833) directly to the plates without changing the medium. The cells were fed daily and passaged as necessary maintaining library coverage until harvest. Cells were harvested for sorting 14 d after transduction of the gRNA sub-library. Cells were washed once with 1x PBS, dissociated using Accutase, filtered through a 30 μm CellTrics filter (Sysmex, 04-004-2326) and resuspended in FACS Buffer (0.5% BSA (Sigma, A7906), 2 mM EDTA (Sigma, E7889) in PBS). Before sorting, an aliquot of 0.4 x 10^6^ cells were taken to represent a bulk unsorted population. The highest and lowest 10% of cells were sorted based on GFP expression and 0.4 x 10^6^ cells were sorted into each bin. Sorting was done with a SH800 FACS Cell Sorter (Sony Biotechnology). After sorting, genomic DNA was harvested with the DNeasy Blood and Tissue Kit (Qiagen, 69506).

### Genomic DNA sequencing

The gRNA libraries were amplified from each gDNA sample across 100 μL PCR reactions using Q5 hot start polymerase (NEB, M0493) with 1 μg of gDNA per reaction. The PCR amplification was done according to the manufacturer’s instructions, using 25 cycles at an annealing temperature of 60 °C with the following primers:

Fwd: 5′-AATGATACGGCGACCACCGAGATCTACACAATTTCTTGGG TAGTTTGCAGTT

Rev: 5′-CAAGCAGAAGACGGCATACGAGAT-(6-bp index sequence)-GACTCGGTGCCACTTTTTCAA

The amplified libraries were purified with Agencourt AMPure XP beads (Beckman Coulter, A63881) using double size selection of 0.65× and then 1× the original volume. Each sample was quantified after purification with the Qubit dsDNA High Sensitivity assay kit (Thermo Fisher, Q32854). Samples were pooled and sequenced on a MiSeq (Illumina) with 21-bp paired-end sequencing using the following custom read and index primers:

Read1: 5′-GATTTCTTGGCTTTATATATCTTGTGGAAAGGACGAAA CACCG Index: 5′-GCTAGTCCGTTATCAACTTGAAAAAGTGGCACCGAGTC

Read2: 5’ - GTTGATAACGGACTAGCCTTATTTAAACTTGCTATGCTGTTTCCAGCATAGCTCTTAAAC

### Data processing and enrichment analysis

FASTQ files were aligned to custom indexes (generated from the bowtie2-build function) using Bowtie 2 (Langmead and Salzberg, 2012). Counts for each gRNA were extracted and used for further analysis. All enrichment analysis was done with R. Individual gRNA enrichment was determined using the DESeq2 (Love et al., 2014) package to compare between high and low, unsorted and low, or unsorted and high conditions for each screen.

### gRNA validations

The top enriched gRNAs from the screens were individually cloned into the appropriate gRNA expression vector as described previously. The gRNA validations were performed similarly as done with the screens using maternal or paternal *SNRPN-2A-GFP* lines stably expressing either dCas9^KRAB^ or ^VP64^dCasd9^VP64^, except the transductions were performed in 24-well plates and the virus was delivered at high MOI. For the validations of the dCas9^KRAB^ paternal screen, single gRNAs were tested per region. For the validations of the ^VP64^dCas9^VP64^ maternal screen, pools of 3-4 gRNAs were tested per region in SNRPN-2A-GFP iPSCs. For validations of the ^Tet1c^dCas9 screen and all studies in WT and ΔPWS iPSCs, single gRNAs were tested except when otherwise specified. Cells were cultured on matrigel-coated 24-well plates in mTesR and harvested for flow cytometry or qRT-PCR 5 -7 d after gRNA transduction for dCas9^KRAB^ and 9-15 days after gRNA transduction for ^VP64^dCas9^VP64^ and ^Tet1c^dCas9. All gRNAs used in this study can be found in Supplementary Table 1.

### Bisulphite primer design, bisulphite conversion, and sequencing

Primer pairs for bisulphite sequencing were designed for the target region using Zymo Research Bisulfite Primer Seeker (<https://www.zymoresearch.com/pages/bisulfite-primer-seeker>), with a maximum amplicon length of 400bp. Each primer set was initially tested on WT and ΔPWS iPSC ^Tet1c^dCas9 stable lines (with no gRNA) with 400ng input gDNA per conversion reaction. To validate primer pair on bisulphite-converted gDNA, we used bisulphite-converted product as input in PCR reactions and tested an annealing temperature gradient ranging from 55 to 62C for 30 cycles with Kapa Uracil+ HotStart ReadyMix polymerase (Roche, 7959052001). Samples of each PCR product were run on a 2% agarose gel. Products displaying bands of the expected size without primer dimer or off-target were Sanger-sequenced (Genewiz) to verify that the correct region was amplified and that the expected proportion of T/C reads were present at each differentially methylated cytosine within the region. Of 4 tested primer pairs, 2 pairs were found to amplify the expected region within the PWS-IC specifically. We proceeded with the primer pair that produces a 400bp amplicon covering 24 CpG sites within the PWS-IC, with the following sequences:

Forward primer 5’-*CTTGCTTCCTGGCACGAG*TTTAAAGTTTTTTGTTTTGGAGAATTAG-3’ Reverse primer 5’-*CAGGAAACAGCTATGAC*CAATAATCACTATTATACACCTACCTAC-3’

In the primers above, italicized sequences are tag sequences that allow for a second round of PCR, and non-italicized sequences bind to the gDNA.

For bisulphite sequencing of iPSCs and neurons, genomic DNA was extracted from cultured cells with the Qiagen DNEasy Blood and Tissue kit (Qiagen, 69504). gDNA was stored at -80C. 250ng of gDNA were input into the bisulphite conversion reaction. For bisulphite conversion and purification, we used the Zymo EZ DNA Methylation Gold kit as instructed (Zymo, D5005). 2uL of bisulphite-converted gDNA was used as input for PCR as described above, with an annealing temperature of 57C. PCR1 reactions were then cleaned with Ampure XP beads (Beckman, A63881) at a ratio of 1.8x. 1/10th of PCR1 was used as input for PCR2 with Q5 polymerase. PCR2 added i5 and i7 barcodes and P5 and P7 overhangs for dual-index Illumina short read sequencing. PCR2 products were purified with Ampure XP beads. Products were visualized via electrophoresis on an Agilent TapeStation 4200 and quantified with Qubit dsDNA HS assay kit (Invitrogen, Q32851) on a Qubit fluorometer (Invitrogen). Libraries were pooled and sequenced on an Illumina Miseq instrument with a Miseq v3 600 cycle kit (Illumina, MS-102-3003) with read lengths of 250x250.

### Bisulphite sequencing data analysis

From the raw read files, we used Trimmomatic (Bolger et al., 2014) to trim reads using the following settings:

HEADCROP: 25; ILLUMINACLIP:TruSeq3-PE-2.fa:2:30:10; TRAILING: 20; SLIDINGWINDOW:4:15; MINLEN:40

Bismark (Krueger and Andrews, 2011) was used to create a bisulphite-converted reference genome from the hg19 genome build. Trimmed, paired-end reads were aligned to the genome and analysed with Bismark version 0.22.3. The output coverage files were used to determine percentage of methylated cytosine for each CpG site in the amplicon.

### HCR FlowFISH

For HCR FlowFISH, reagents and probe sets were ordered from Molecular Instruments (<https://www.molecularinstruments.com/>). Buffer set formulations were for cells in suspension. Approximately 2 million cells per sample were dissociated from the plate with Accutase and processed according to the protocol as described in Reilly et al. 2021, except that probe hybridisation buffer, probe wash buffer, and amplification buffer were used from Molecular Instruments (Reilly et al., 2021){Krueger 2011}. Probe sets were used at 4nM concentration. Fixation and permeabilisation buffer was prepared fresh as 4% paraformaldehyde (Electron Microscopy Sciences, 15710) and 0.1% Tween-20 (Roche, 11332465001) in 1x divalent cation-free phosphate-buffered saline. After staining, cells were analysed on a Sony SH800Z cell sorter.

### ATAC sequencing

iPSCs were dissociated from the plate using Accutase, and viability and cell number were assessed with a Countess II cell counter (Invitrogen) and Trypan Blue (Invitrogen, T10282). 45,000 cells per sample were processed for ATAC-seq according to the Omni-ATAC protocol (Corces et al., 2017). Libraries were sequenced on an Illumina NovaSeq with 2x25 paired-end reads with a coverage of 50 million reads per sample. Reads were trimmed with Trimmomatic (Bolger et al., 2014) to remove adapter content and filter on quality score, then trimmed reads were aligned to hg19 using Bowtie2 (Langmead and Salzberg, 2012). Reads mapping to ENCODE blacklisted regions were removed, and duplicated reads were removed with Picard MarkDuplicates. Peaks were called using MACS2 (Zhang et al., 2008). For visualization, bamCoverage was used to generate rpkm-normalized bigwig files from deduplicated bam files. Quality of each sample was assessed based on number of uniquely mapping reads after blacklist removal. We generated a union peak set from narrowPeak files of all samples. Counts files for each sample were generated using the featureCounts function. Differential peak analysis was conducted with DESeq2 on the feature counts files with an adjusted p-value threshold of p<0.01 for differential peak analysis.

---

**Supplementary Figure S1, related to Figure 1.**
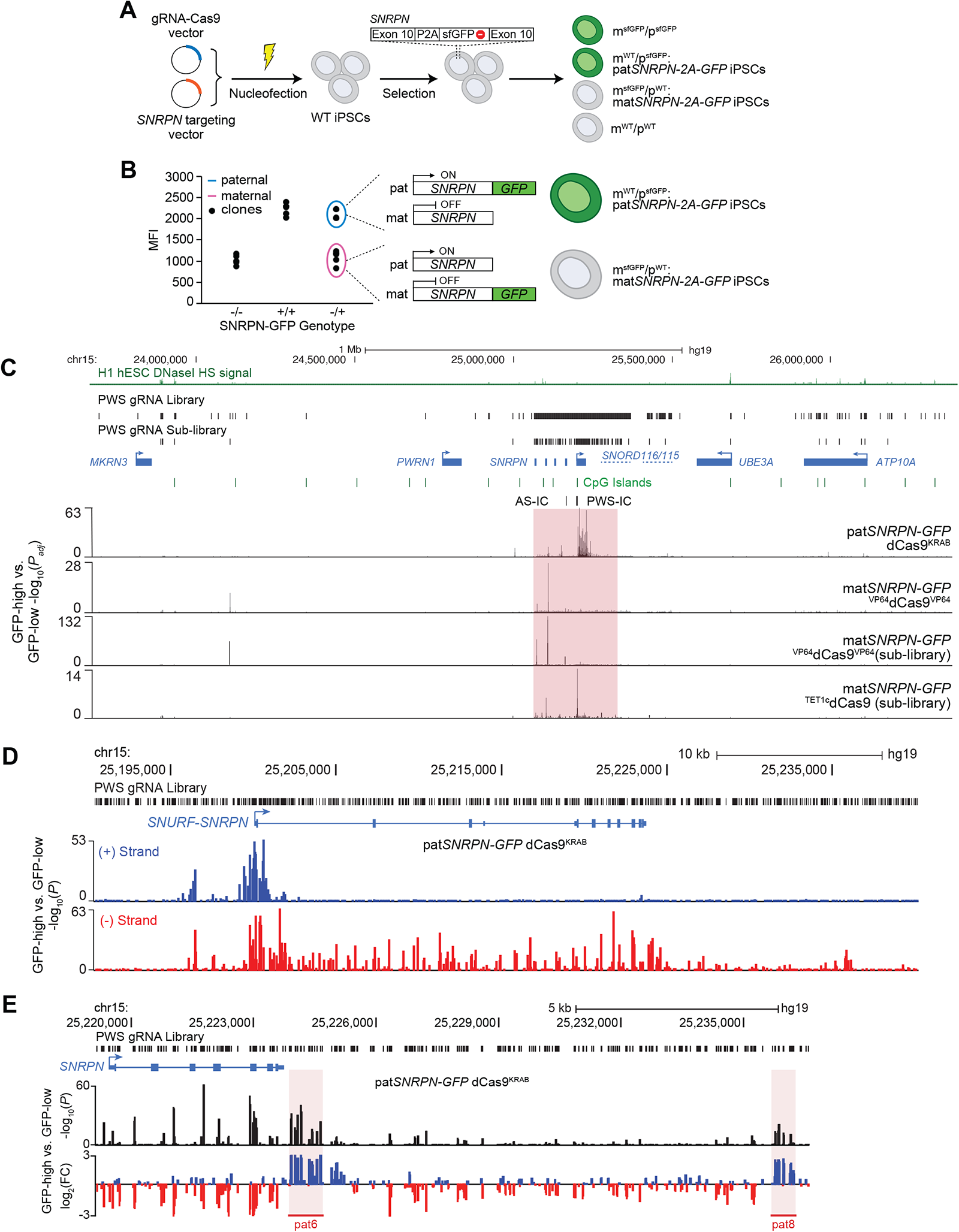
CRISPRa/i screens. (A) Schematic of derivation of maternally or paternally tagged *SNRPN-2A-GFP* iPSCs. (B) Flow cytometry validation of differential GFP fluorescence of maternally and paternally-tagged lines. (C) CRISPR screen results shown in Fig. 1D, view zoomed out to cover the entire span of the human PWS gRNA library. The red highlight indicates the area pictured in Fig. 1D. (D) CRISPRi dCas9^KRAB^ *patSNRPN-2A-GFP* screen results (shown in Fig. 1(D)), separated by strand on which the gRNA is located. (E) CRISPRi screen results at the 3’ end of *SNRPN* (as shown in Fig. 1(D)), plotted as log_2_(fold change) gRNA enrichment between low and high GFP sorted bins.

**Supplementary Figure S2, related to Figure 1.**
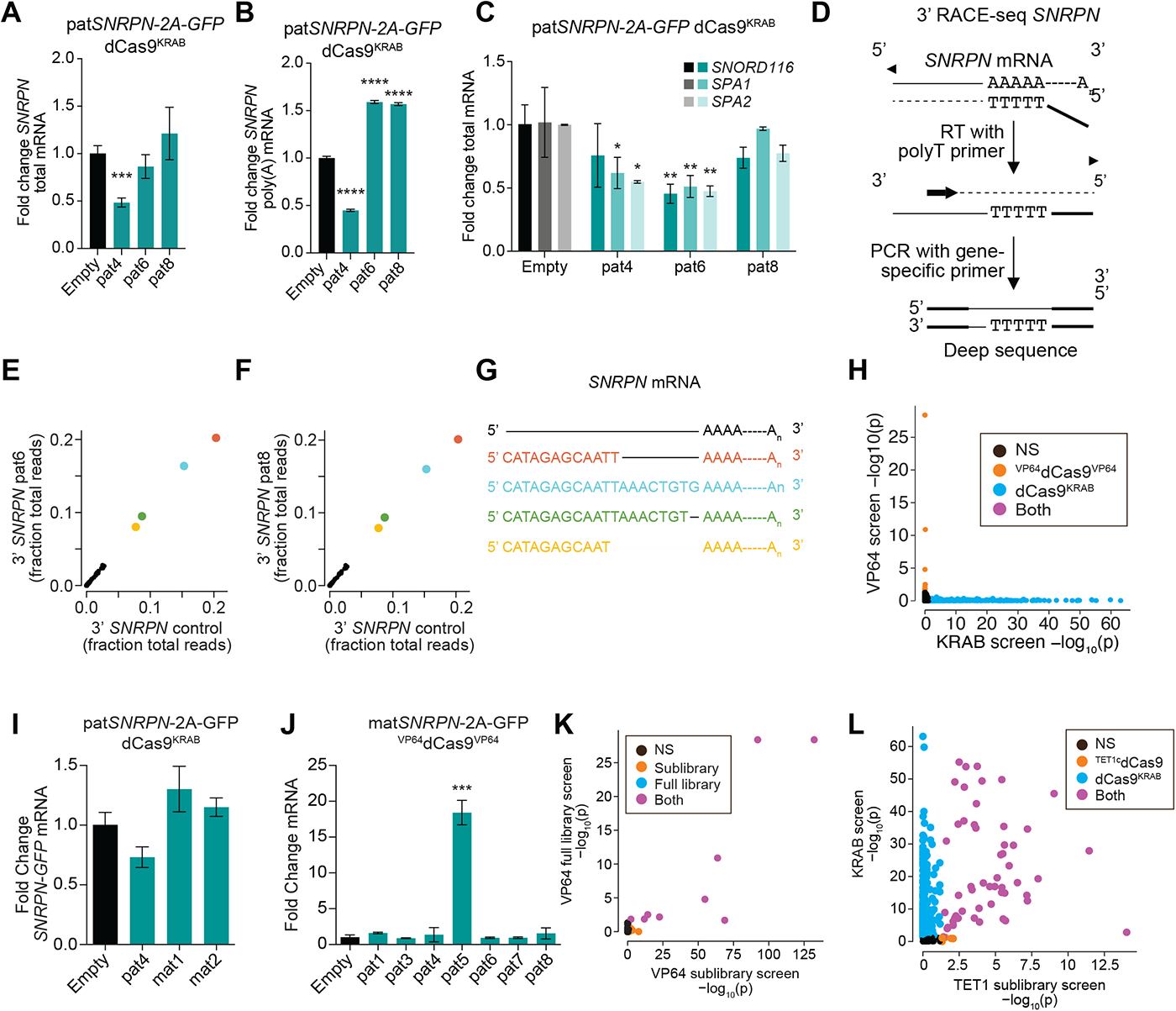
Further validations of CRISPRa/i screens. (A) qPCR of *SNRPN-GFP* from total mRNA in *patSNRPN-2A-GFP* dCas9^KRAB^ lines. (B) qPCR of *SNRPN-GFP* from polyadenylated (poly-A) mRNA in *patSNRPN-2A-GFP* dCas9^KRAB^ lines. (C) qPCR of the indicated genes from total mRNA in *patSNRPN-2A-GFP* dCas9^KRAB^ lines. (D) Schematic of 3’ RACE-seq of *SNRPN* transcript. (E,F) Comparison of *SNRPN* 3’ UTR sequence variants in control cells with dCas9^KRAB^ and an empty gRNA vector and cells treated with either (E) a pat6 gRNA or (F) a pat8 gRNA (G) Sequences of the four most predominant 3’ UTR variants detected in all conditions. Colors in (E) and (F) match the correspondingly colored sequences in (G). (H) Plot of -log_10_(padj) values of each gRNA in the ^VP64^dCas9^VP64^ full library screen vs. dCas9^KRAB^ screen. (Significant p_adj_ < 0.05.) (I) qPCR of *SNRPN-GFP* in *patSNRPN-2A-GFP* dCas9^KRAB^ iPSCs with gRNAs from the pat4, mat1, and mat2 regions. (J) qPCR of *SNRPN-GFP* in *matSNRPN-2A-GFP* ^VP64^dCas9^VP64^ iPSCs with gRNAs from the CRISPRi screen hits. (K) Plot of -log_10_(p_adj_) values of each gRNA in the ^VP64^dCas9^VP64^ full library screen vs. sublibrary screen. (Significant p_adj_ < 0.05.) (L) Plot of -log_10_(p_adj_) values of each gRNA in the ^Tet1c^dCas9 sublibrary screen vs. dCas9^KRAB^ full library screen. Significant p_adj_ < 0.05.

**Supplementary Figure S3, related to Figures 2 and 3.**
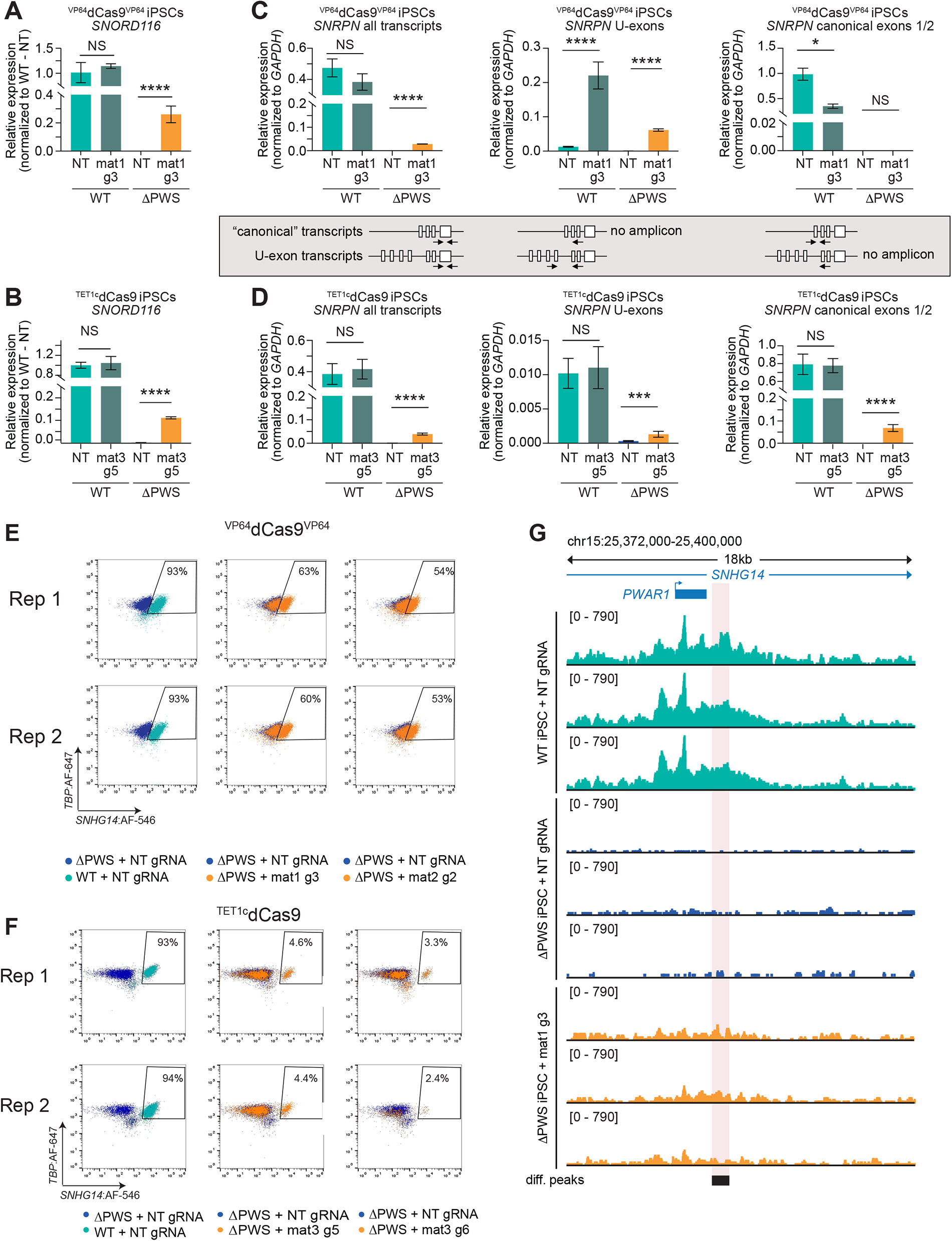
PWS gene expression in ^VP64^dCas9^VP64^ or ^Tet1c^dCas9 ΔPWS iPSCs. (A,C): qPCR of WT or ΔPWS ^VP64^dCas9^VP64^ iPSCs with NT or mat1 g3 gRNA for either (A) *SNORD116* or (C) sets of *SNRPN* transcript variants. (B,D) qPCR of WT or ΔPWS ^Tet1c^dCas9 iPSCs with NT or mat3 g5 gRNA for either (B) *SNORD116* or (D) sets of *SNRPN* transcript variants. Fold change values (relative to *GAPDH*) are plotted mean +/-SD, but statistics were calculated on dCt values (normalized to *GAPDH*); one-way ANOVA followed by Sidak’s multiple comparisons test for select groups WT + targeting gRNA vs. NT gRNA or ΔPWS + targeting gRNA vs. NT gRNA. *p<0.05, ***p<0.001,****p<0.0001. (E,F) Two replicates of HCR FlowFISH (Rep. 1 of each was shown in Fig. 2F,G, respectively) of WT or ΔPWS iPSCs with either (E) ^VP64^dCas9^VP64^ and NT or mat1/2 gRNAs, or (F) ^Tet1c^dCas9 and NT or mat3 gRNAs. (G) Browser tracks of ATAC sequencing (rpkm-normalized BigWig) of ΔPWS or WT iPSCs with ^VP64^dCas9^VP64^ and NT or mat1 g3 gRNA downstream of *SNRPN,* at the *PWAR1* gene.

**Supplementary Figure S4, related to Figures 2 and 3.**
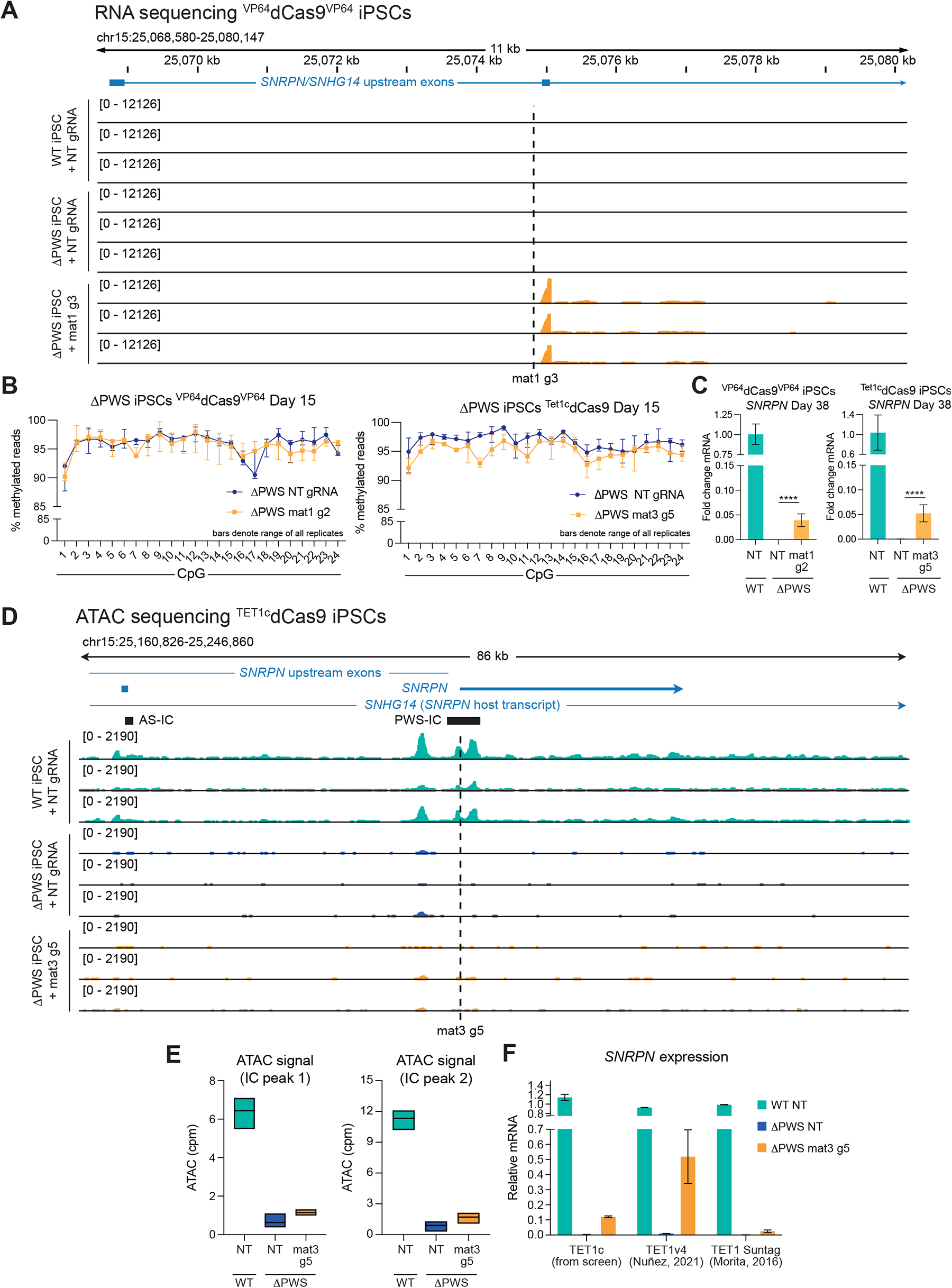
Additional sequencing of ^VP64^dCas9^VP64^ or ^Tet1c^dCas9 ΔPWS iPSC conditions. (A) Genome browser visualization of RNA sequencing (rpkm-normalized BigWig) of ^VP64^dCas9^VP64^ WT or ΔPWS iPSCs with NT or mat1 g3 gRNA, zoomed in on *SNRPN* upstream exons. (B) Targeted bisulphite sequencing of ΔPWS iPSCs with ^VP64^dCas9^VP64^ or ^Tet1c^dCas9 covering 24 CpG sites within the PWS locus (hg19 chr15: 25200353-25200693), 15 days post-transduction. Data are shown as the range of the data, with the plotted point being the median, n = 3 replicates. (C) qPCR of *SNRPN* in WT or ΔPWS iPSCs with ^VP64^dCas9^VP64^ or ^Tet1c^dCas9 38 days after transduction with the indicated gRNA, same cell samples as those collected for bisulphite sequencing depicted in Figure 3A,B. For both qPCR plots, fold change values are plotted mean +/-SD, but statistics were calculated on ddCt values (normalized to *GAPDH* and WT ctrl sample); one-way ANOVA followed by Dunnett’s test vs. ΔPWS NT gRNA ****p<0.0001. (D) Browser tracks of ATAC sequencing (rpkm-normalized BigWig) of ΔPWS or WT iPSCs with ^Tet1c^dCas9 and NT or mat3 g5 gRNA. (E) Quantification of ATAC-seq reads (counts per million) at each of the two peaks at the PWS-IC (mat3 g5 is located within the first of the two peaks, see S4D). ΔPWS + NT vs. mat3 gRNA not significant, Tukey’s test following one-way ANOVA. (F) qPCR of SNRPN expression in ΔPWS iPSCs with NT or mat3 g5 gRNA, comparing 3 different ^Tet1c^dCas9 constructs, all delivered by lentivirus.

